# Design Automation of Microfluidic Single and Double Emulsion Droplets with Machine Learning

**DOI:** 10.1101/2023.05.31.543128

**Authors:** Ali Lashkaripour, David P. McIntyre, Suzanne G.K. Calhoun, Karl Krauth, Douglas M. Densmore, Polly M. Fordyce

**Affiliations:** Department of Bioengineering, Stanford University, Stanford, CA, USA; Department of Genetics, Stanford University, Stanford, CA, USA; Biomedical Engineering Department, Boston University, MA, USA; Biological Design Center, Boston University, Boston, MA, USA; Chemical Engineering Department, Stanford University, Stanford, CA, USA; Electrical & Computer Engineering Department, Boston University, Boston, MA, USA; Chan-Zuckerberg Biohub, San Francisco, CA, USA

## Abstract

Droplet microfluidics enables kHz screening of picoliter samples at a fraction of the cost of other high-throughput approaches. However, generating stable droplets with desired characteristics typically requires labor-intensive empirical optimization of device designs and flow conditions *that limit adoption to specialist labs*. Here, we compile the most comprehensive droplet dataset to date and use it to train machine learning models capable of accurately predicting device geometries and flow conditions required to generate stable aqueous-in-oil and oil-in-aqueous single and double emulsions from 15 to 250 *μ*m at rates up to 12000 Hz for different fluids commonly used in life sciences. Novel device geometries predicted by our models for as-yet-unseen fluids yield accurate predictions, establishing their generalizability. Finally, we generate an easy-to-use design automation tool that yield droplets within 3 *μ*m (< 8%) of the desired diameter, facilitating tailored droplet-based platforms for new applications and accelerating their utility in life sciences.

## 1 Introduction

Droplet microfluidics enables massively parallel miniaturized assays by stably dispersing nanoliter to picoliter samples of a liquid (the ‘dispersed’ fluid) within an immiscible carrier liquid (the ‘continuous’ fluid) [1]. Single emulsion (SE) water-in-oil or oil-in-water droplet systems have unlocked new opportunities in single-cell omics [2–4], directed evolution [5, 6], chemical synthesis [7], and drug and antibody discovery [8, 9]. Double emulsion (DE) droplets commonly consist of an aqueous core wrapped in an oil shell that is dispersed in an aqueous outer continuous fluid [10] and have been used for controlled drug delivery [11, 12], production of microparticles with core–shell structures [13, 14], and in the food and cosmetics industry [15, 16]. Due to their aqueous outer fluid and high stability, DEs can also be sorted using commercial fluorescence-activated cell sorting (FACS) machines, enabling off-the-shelf screening in droplet microfluidics at kHz throughput [17].

Despite the benefits of droplet microfluidics, adoption of this technology in life sciences has been limited primarily to specialized groups or commercially available products with limited functionality (e.g., 10x Genomics Chromium machines [18, 19]), largely because droplet stability, size, and generation rates dictate downstream assay performance but are difficult to predict. The effective concentration rates of species in droplet assays scales inversely with the 3^*rd*^ power of droplet diameter, and single-cell encapsulation and the efficiency of FACS sorting are highly size-dependent [20, 21]. Precise control over generation rate is similarly crucial for development of multi-component microfluidic platforms [22].

Droplets are most commonly made using flow-focusing geometries that yield highly monodisperse droplets over a wide range of diameters and generation rates and require lower continuous-to-dispersed flow rate ratios [23–26]. However, the complex and highly nonlinear dynamics of multiphase flows and a large number of effective parameters in flow-focusing geometries have made it difficult to establish an analytical solution or a generalizable scaling formula that can accurately predict droplet diameter and rate across a broad range of flow conditions and fluid properties [27, 28]. These limitations are exacerbated in life sciences given that biological assays require buffers with varying properties (e.g., interfacial tension and viscosity) that can significantly impact the resultant droplet diameter and generation rate [29, 30]. As a result, generating droplets with desired properties typically requires multiple resource-intensive design iterations and empirical tests [31, 32], and this process becomes even more challenging when integrating other components upstream or downstream of a droplet generator [22, 33]. Thus, a predictive understanding of droplet generation would enable conversion of high-level performance requirements to a microfluidic device design, facilitate multi-component devices, and facilitate broader adoption of these platforms in life sciences [32].

Machine learning models trained on experimental data were recently demonstrated to enable accurate prediction of SE droplet generation performance [34]. However, previously proposed models only account for variations in flow rates and device geometries [35] or surfactants [36]. As a result, previous models offer limited utility in life science applications. Here, we leverage machine learning and a comprehensive experimental dataset including both SE and DE droplets comprised of many different fluids to train models that accurately predict droplet diameter and generation rate across a diverse range of fluid properties, geometries, flow rates, and device materials. Additionally, we demonstrate that our models generalize to new geometries and fluids by experimentally validating ‘blind’ predictions using novel device geometries and fluid compositions. Finally, we integrate these predictive models with an automated search algorithm to create a design automation tool for SE and DE droplets. This opensource tool, called DAFD 3.0 (Design Automation of Fluid Dynamics), can return the necessary design and flow rates to achieve the user-specified diameter and rate for different fluids, while also predicting other characteristics such as performance range and stability (Fig. 1).

**Figure 1:**
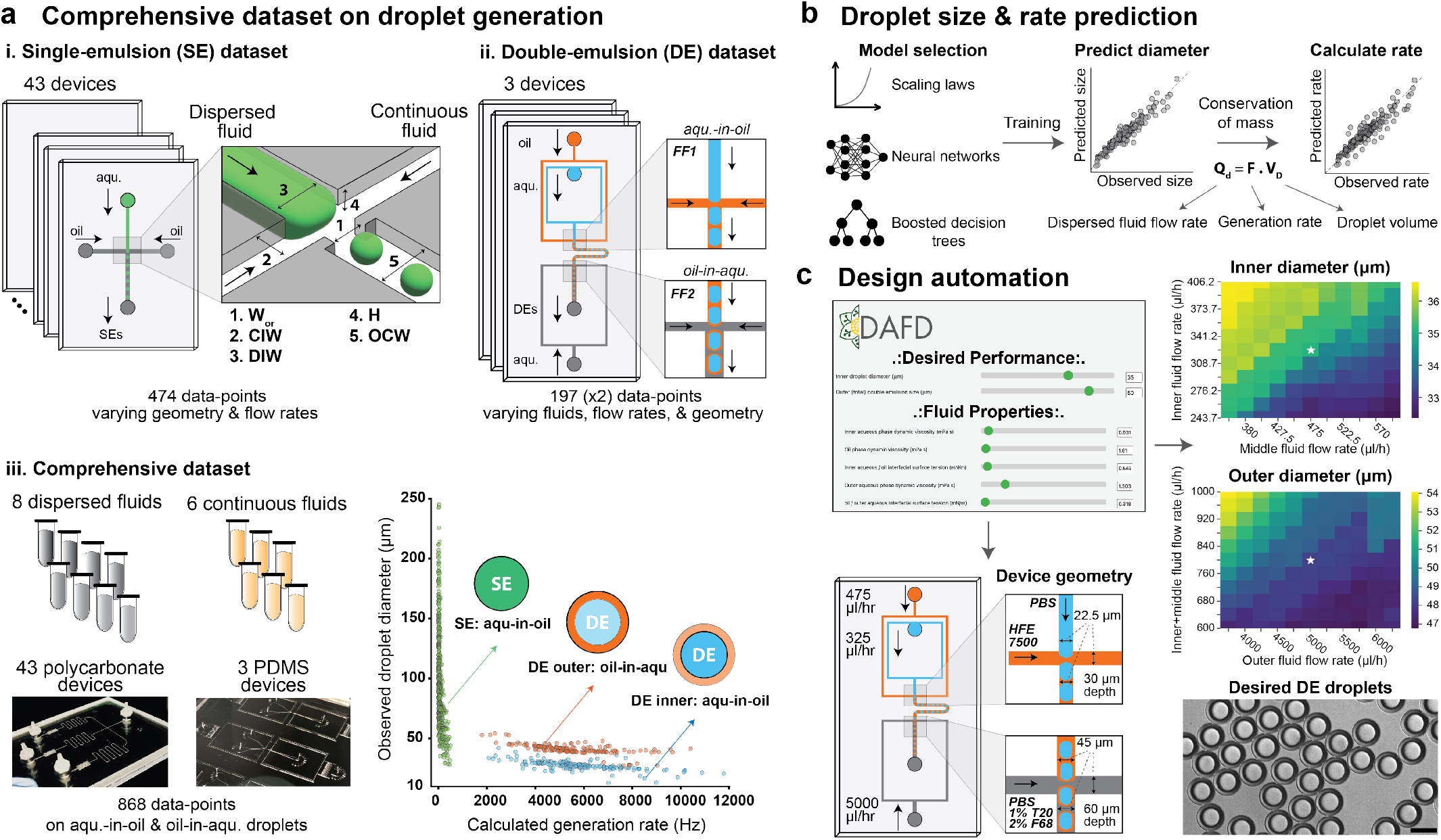
Pipeline for collating data and training models to enable performance prediction and design automation of SE and DE droplet generation. **a**. Composition of datasets exploring effects of geometry, fluid properties, and flow rates on (i.) SE and (ii.) DE droplet generation collated to yield a final (iii.) comprehensive dataset with 868 entries. The combined dataset includes 8 dispersed fluids, 6 continuous fluids, and 46 devices that yield aqueous-in-oil and oil-in-aqueous droplets with diameters from 15 to 250 *μ*m at rates of 5 to 12000 Hz. **b**. Schematic of model training to predict: (1) droplet diameter based on device geometry, fluid properties, and flow rates, and (2) droplet generation rates based on predicted diameters and conservation of mass (see Methods). **c**. Predictive models were integrated with a custom search algorithm to convert user-specified desired droplet characteristics to an optional device design and flow rates. This open-source software tool, DAFD 3.0, is available at: dafdcad.org.

## 2 Results

### 2.1 Comprehensive droplet generation dataset

To generate a comprehensive dataset detailing impacts of device designs, flow rates, and fluid properties (e.g., viscosity and interfacial tension) on droplet diameters and generation rates, we curated and combined two previously generated SE and DE experimental datasets [30, 35]. This comprehensive dataset includes 46 different polydimethylsiloxane (PDMS) and polycarbonate device designs (43 SE and 3 DE generators with 49 flow-focusing geometries combined), 8 different dispersed fluids, and 6 different continuous fluids for generating aqueous-in-oil and oil-in-aqueous droplets of 15 to 250 *μ*m in diameter at rates of 5 to 12000 Hz (Fig. 2).

**Figure 2:**
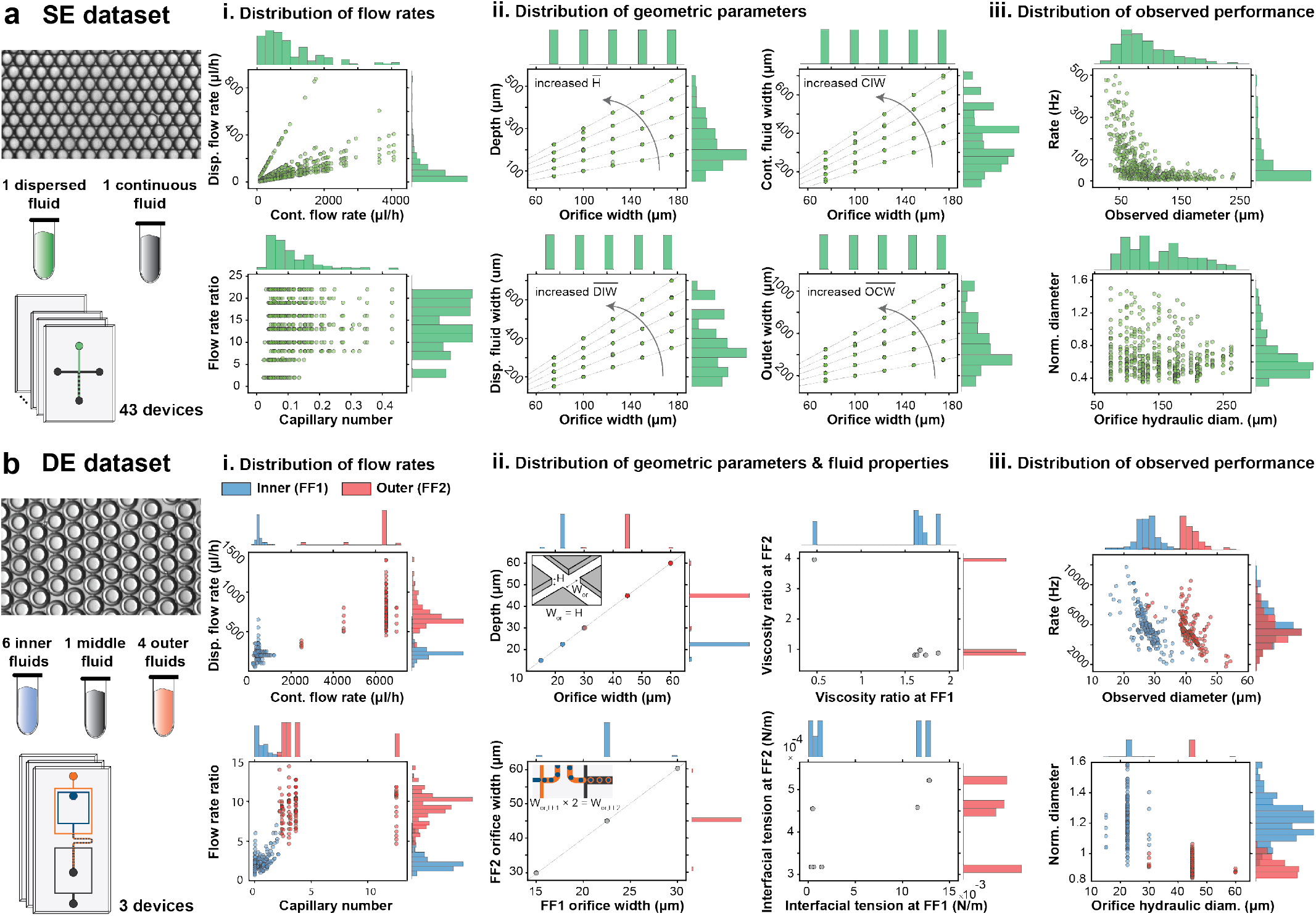
The comprehensive dataset includes SE and DE droplets produced using a wide variety of device geometries, fluid properties, flow rates. **a**. The SE dataset includes 43 polycarbonate devices that varied orifice width, channel depth, outlet width, and continuous and dispersed fluid inlet widths; each device was used over multiple flow rate ratios and capillary numbers to generate SEs with diameters of 25 to 250 *μ*m at rates of 5 to 500 Hz. **b**. The DE dataset includes 3 PDMS devices (each with two flow-focusers) and 6 inner, 1 middle, and 4 outer fluids. Devices were tested at different flow rates of inner, oil, and outer fluids (different capillary numbers and flow rate ratios for each junction). The orifice widths of the PDMS devices were 15, 22.5, and 30 *μ*m at the first junction, with an aspect ratio of 1, while the orifice size at the 2^*nd*^ junction was twice the size of the 1^*st*^ junction. Several biologically relevant fluids with different viscosities and interfacial tensions were used to generate droplets with diameters of 15 (inner) to 54.2 (outer) *μ*m at rates of 1800 to 11800 Hz.

#### 2.1.1 Single emulsion droplets

We previously generated aqueous-in-oil (DI water and mineral oil) SEs using 43 devices and multiple capillary numbers and flow rate ratios (see Methods for definitions) [35]. This dataset varies the orifice width from 75 to 175 *μ*m and systematically explores the remaining geometric parameters according to the orifice width (Fig. 2a). The devices were then tested at a range of capillary numbers and flow rate ratios and yielded droplets of 25 to 250 *μ*m at 5 to 500 Hz in the dripping regime (474 datapoints total). To improve generalizability, we first converted the orifice dimensions of each device to hydraulic diameter (*D*_*h*_):

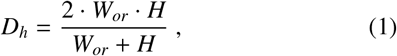

where *W*_*or*_ is orifice width and *H* is channel height. We then computed normalized droplet diameter 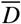 produced by each device:

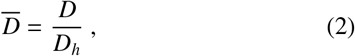

where *D* is the observed droplet diameter. The diverse range of flow rates (Fig. 2a.i) and device design parameters (Fig. 2a.ii) resulted in droplets with normalized diameters ranging from 0.35 to 1.5 (Fig. 2a.iii).

#### 2.1.2 Double emulsion droplets

We also previously generated aqueous-oil-aqueous DEs using 3 PDMS devices with different geometries using multiple flow rates (197 datapoints total). These experiments used several biologically relevant fluids with applications in cell culture, cell lysis, and molecular biology (e.g., PCR, NGS, and ATAC-Seq) including 6 different inner, 1 middle, and 4 outer fluids (Table 1) [30]. The 3 devices contained orifice widths of 15, 22.5, and 30 *μ*m at flow-focuser 1 (FF1) and 30, 45, and 60 *μ*m at flow-focuser 2 (FF2), respectively, with a normalized depth (i.e., aspect ratio) of 1. The orifice at FF2 is twice as wide and deep as orifice at FF1. The resultant droplet diameters ranged from 15.5 *μ*m to 54.2 *μ*m and generation rates varied from 1800 to 11800 Hz. To effectively model DE generation, we considered it as two independent SE generation events at FF1 and FF2, with FF1 generating aqueous-in-oil SEs and FF2 generating oil-in-aqueous SEs (we validate this assumption in Performance prediction section). We also normalized DE inner and outer diameters using the hydraulic diameters at FF1 and FF2, respectively (i.e., the orifice at which droplets are generated). Normalized inner diameters varied from 0.92 to 1.6 (15.5 to 42.1 *μ*m) and normalized outer diameters ranged from 0.84 to 1.06 (27.4 to 54.2 *μ*m) as shown in Fig. 2b.

**Table 1:** 11 different fluid combinations in the dataset make it possible to investigate effects of fluid properties on droplet generation. These fluids are commonly used across different life science applications. The viscosity of the dispersed fluids varied from 0.86 to 3.4 mPa.s and the viscosity of the continuous fluids varied from 1.61 to 57.2 mPa.s. The interfacial tension between the dispersed and continuous fluids ranged from 0.318 to 12.84 mN/m. Newly generated data and previously published data on droplet generation using 4 new and unseen fluid combinations were used to assess the generalizability of models to novel fluids.

| Dispersed fluid |  | Continuous fluid |  | Interaction |
| --- | --- | --- | --- | --- |
| Fluid | Viscosity (mPa.s) | Fluid | Viscosity (mPa.s) | Interfacial tension (mN/m) |
| **Fluids included in the comprehensive dataset** |  |  |  |  |
| 1. M9 bacterial media | 0.861 | HFE 7500 oil + 2.2% ionic Krytox | 1.61 | 12.84 |
| 2. M9 bacterial media + 25 mM glucose | 0.967 | HFE 7500 oil + 2.2% ionic Krytox | 1.61 | 11.60 |
| 3. PBS | 0.931 | HFE 7500 oil + 2.2% ionic Krytox | 1.61 | 0.543 |
| 4. PBS + 1% Tween-20 | 0.988 | HFE 7500 oil + 2.2% ionic Krytox | 1.61 | 0.319 |
| 5. PBS + 0.9% NP40 | 1.003 | HFE 7500 oil + 2.2% ionic Krytox | 1.61 | 1.41 |
| 6. PBS + 10% PEG 6000 mw | 3.431 | HFE 7500 oil + 2.2% ionic Krytox | 1.61 | 0.461 |
| 7. DI water | 1.001 | NF350 mineral oil + 5% Span-80 | 57.2 | 5.0 |
| 8. HFE 7500 oil + 2.2% ionic Krytox | 1.61 | PBS+1% Tween-20 + 2% Pluronic F68 | 1.303 | 0.318 |
| 9. HFE 7500 oil + 2.2% ionic Krytox | 1.61 | M9 salts + 2% Pluronic F68 | 1.412 | 0.522 |
| 10. HFE 7500 oil + 2.2% ionic Krytox | 1.61 | M9 salts + 25 mM glucose + 2% Pluronic F68 | 1.563 | 0.458 |
| 11. HFE 7500 oil + 2.2% ionic Krytox | 1.61 | PBS + 10% PEG 6000mw + 2% Pluronic F68 | 6.395 | 0.455 |
| **Unseen fluids for assessing the generalizability of models** |  |  |  |  |
| 12. DMEM complete cell media + 16% Optiprep | 1.25 | 2% dSurf HFE 7500 oil | 1.61 | 9.74 |
| 13. RPMI 1640 complete cell media + 20% Optiprep + 0.1% Pluronic F127 | 1.36 | 2% dSurf HFE 7500 oil | 1.61 | 6.16 |
| 14. 2% dSurf HFE 7500 oil | 1.61 | RPMI 1640 complete cell media + 5% Pluronic F127 | 2.82 | 4.61 |
| 15. Surfactant-free HFE 7500 oil | 1.31 | trimethylolpropane trimethacrylate (TMPTMA) | 4.2 | 3.10 |

We then curated and combined the SE and DE datasets by using standardized definitions of capillary number and geometric parameters to create a comprehensive dataset of microfluidic droplet generation that covers a diverse design space of capillary numbers, flow rate ratios (Supplementary Fig. 1), geometries, fluid properties, flow conditions, and output performance (Table 2). As orifice length minimally impacts droplet generation in the dripping regime, we did not consider it as a design parameter [37]. This enabled us to model flow-focusing geometries that do not contain an orifice constriction, where orifice length cannot be clearly defined. This dataset is the largest experimental dataset available for microfluidic droplet generation in the dripping regime and is the first to include aqueous-in-oil and oil-in-aqueous droplets, different biologically relevant fluids and device materials (Supplementary Table 1).

**Table 2:** The comprehensive dataset includes 868 datapoints on single and double emulsion generation with different fluids. This dataset includes both aqueous-in-oil and oil-in-aqueous droplets with a broad range of output performance. This is achieved by varying effective parameters in flow-focusing droplet generation including device geometry, fluid properties, and flow rates.

| Parameter | Unit | Lower bound | Upper bound | Unique values |
| --- | --- | --- | --- | --- |
| **Output performance** |  |  |  |  |
| Droplet diameter | $\mu\text{m}$ | 15.5 | 245.1 | 825 |
| Generation rate | Hz | 5 | 11,774 | 838 |
| Droplet diameter normalized by hydraulic diameter | N.A. | 0.35 | 1.60 | 833 |
| **Fluid properties** |  |  |  |  |
| Dispersed fluid viscosity | mPa.s | 0.861 | 3.431 | 8 |
| Continuous fluid viscosity | mPa.s | 1.303 | 57.2 | 6 |
| Interfacial tension | mN.m<sup>-1</sup> | 0.318 | 12.84 | 11 |
| Viscosity ratio (continuous/dispersed) | N.A. | 0.47 | 57.2 | 11 |
| **Geometric parameters** |  |  |  |  |
| Orifice width | $\mu\text{m}$ | 15 | 175 | 10 |
| Normalized channel depth | N.A. | 1 | 3 | 13 |
| Normalized continuous fluid inlet width | N.A. | 1 | 4 | 15 |
| Normalized dispersed fluid inlet width | N.A. | 1 | 4 | 13 |
| Normalized outlet channel width | N.A. | 1 | 6 | 15 |
| **Flow parameters** |  |  |  |  |
| Flow rate ratio (continuous/dispersed) | N.A. | 0.69 | 22 | 187 |
| Capillary number | N.A. | 0.014 | 9.399 | 206 |

### 2.2 Droplet diameter and generation rate prediction

We trained scaling law, neural network, and boosted decision tree models to predict SE and DE droplet diameters and generation rates. While scaling laws (i.e., empirically fitted scaling formulas) are simple and have been traditionally used for this task, they are often inaccurate or fail to generalize to unseen fluids and size scales [35, 36]. We therefore also trained machine learning models and compared their accuracy and generalizability to scaling laws. To improve generalizability, we made all design parameters dimensionless when possible. This involved using capillary number, viscosity ratio, and flow rate ratio to account for fluid properties (i.e., viscosity and interfacial tension) and flow rates. We also normalized all geometric parameters (channel depth, dispersed and continuous inlet widths, and outlet channel width) by the orifice width, apart from orifice width itself (Fig. 3a). In all models, we first predicted normalized droplet diameter based on input parameters and then used the hydraulic diameter of the orifice to calculate an actual droplet diameter:

**Figure 3:**
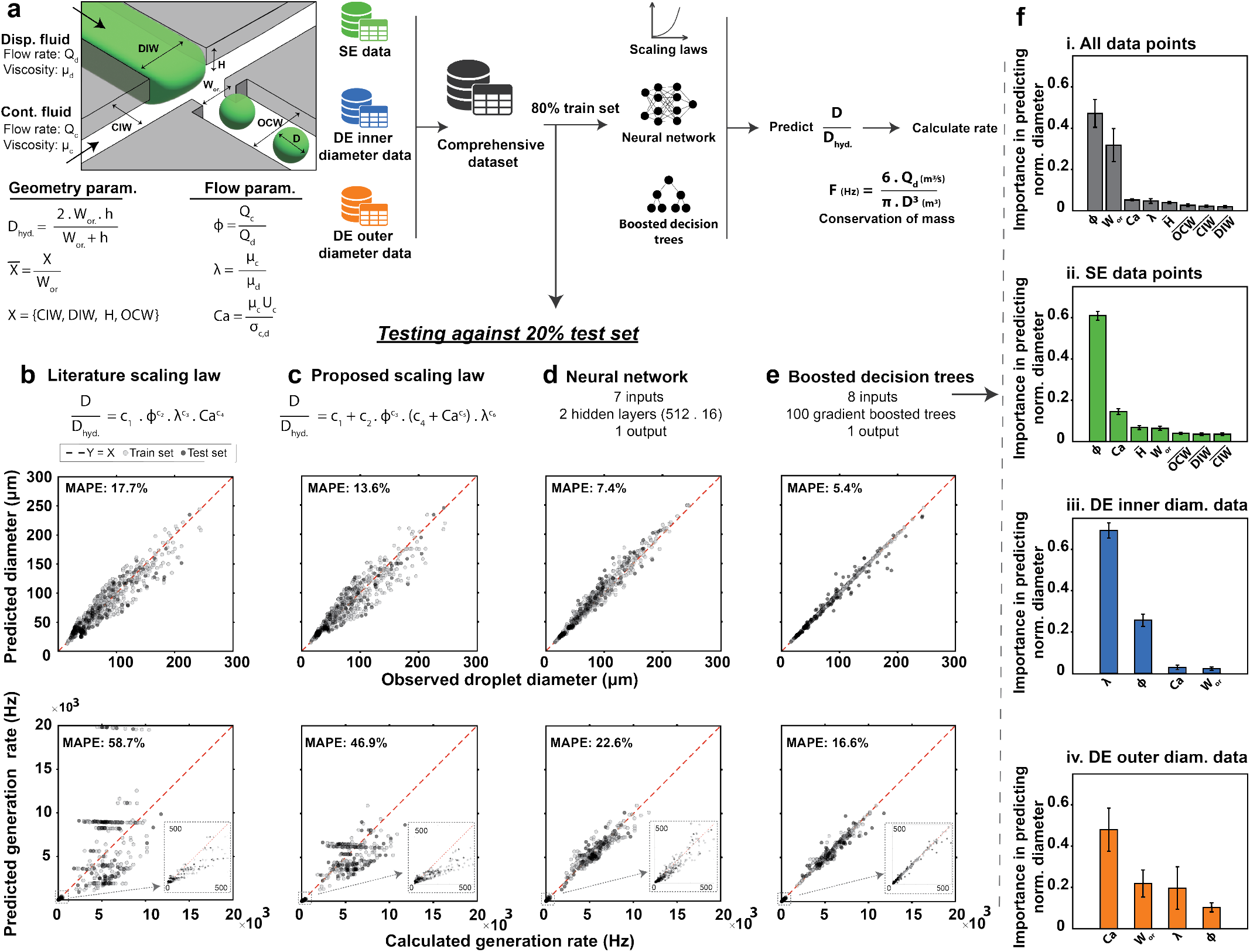
Boosted decision trees and neural networks accurately predict SE and DE droplet diameters and generation rates. **a**. To develop generalizable models, we converted fluid and flow properties and geometric parameters to dimensionless inputs and split the comprehensive dataset into 80% train and 20% test sets across 15 randomized sessions. In each case, we trained and compared performance of **b**. a previously published scaling law [38], **c**. a newly-proposed scaling law, **d**. a neural network and **e**. boosted decision trees. MAPEs for predicting rate were approximately 3 times the MAPEs for diameter, as expected from conservation of mass. Red dashed line indicates the 1:1 line, each grey marker indicates model-predicted values for datapoints included either within the training set (light grey) or the test set (dark grey) of a single representative model. **f**. Relative importance of different parameters in predicting droplet diameters with boosted decision trees; bars represent the average significant and error bars represent standard deviation across 15 random training sessions.

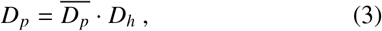

where *D*_*p*_ is predicted droplet diameter, 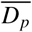 is predicted normalized diameter, and *D*_*h*_ is the hydraulic diameter of the orifice.

To evaluate the accuracy of models and prevent over-fitting, we randomly split the comprehensive dataset into a training set (80%) and a testing set (20%) over 15 different training sessions and calculated the average performance of each model against the test set. For each model, we first predicted droplet diameters and then used these values to calculate predicted droplet generation rates based on dispersed fluid flow rate and conservation of mass (assuming stable droplet generation with a uniform diameter):

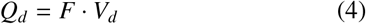

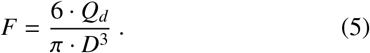

Here, *Q*_*d*_ is the dispersed fluid flow rate, *F* is the generation rate, *V*_*d*_ is the droplet volume, and *D* is the droplet diameter.

To predict DE outer diameters, we set the flow rate of the dispersed fluid to the total flow rate of inner and middle fluids (as is required to satisfy conservation of mass).

#### 2.2.1 Scaling laws

Fitting several previously published scaling laws [38–43] to the comprehensive dataset yielded predictions with a mean absolute percentage error (MAPE) range of 17.7–47.6% for diameter predictions and 58.7–3023% for rate predictions (Supplementary Note 1, Supplementary Table 2 and Supplementary Figs. 2–12). Among these models, the Liu et al. scaling law showed the best accuracy and used flow rate ratio, viscosity ratio, and capillary number as inputs (Fig. 3b) [38]. These inputs may not always affect diameter independently and the impact of flow rate ratio can vary from low to high capillary numbers [37]. Therefore, we also proposed a new scaling law that accounts for some level of parameter dependence. This new scaling law was able to predict diameter and generation rate with a MAPE of 13.6% and 46.9%, respectively (Fig. 3c). Including additional parameters as inputs either prevented finding a solution or reduced accuracy.

#### 2.2.2 Machine learning models

Next, we trained a neural network that takes capillary number, flow rate ratio, and five geometric parameters (orifice width, normalized channel depth, normalized outlet width, normalized dispersed fluid inlet width, and normalized continuous fluid inlet width) as inputs and predicts normalized droplet diameter. We chose a wide and shallow network structure, with 2 hidden layers of 512 and 16 nodes, respectively, which is more suitable for small datasets compared to deep and narrow structures (i.e., more hidden layers with fewer nodes) [44]. The trained neural network significantly outperformed the scaling laws over 15 randomized sessions, with MAPE of 7.4% for diameter and 22.6% for generation rate (Fig. 3d, see Supplementary Fig. 13 for 14 additional training sessions). We did not include viscosity ratio as an input for the neural network as it resulted in a slightly lower accuracy when predicting previously published data not included in the original training and testing datasets (discussed in Generalizability to unseen geometries and fluids section), despite achieving a slightly higher accuracy for the comprehensive dataset (Supplementary Note 2 and Supplementary Table 3).

**Table 3:** Performance prediction accuracy for each model. Metrics are reported for a 20% test-set, using the average plus-minus (±) the standard deviation for 15 different randomized training and testing sessions. ^*^ Mean absolute percentage error, ^**^ Coefficient of determination, ^***^ Mean absolute error, ^****^ Root mean square error.

| Parameter | MAPE * | $R^2$** | MAE*** | RMSE**** |
| --- | --- | --- | --- | --- |
| **Droplet diameter prediction** |  |  |  |  |
| Boosted decision trees | $5.38 \pm 0.34 \%$ | $0.96 \pm 0.00$ | $4.62 \pm 0.41 \mu\text{m}$ | $8.29 \pm 0.73 \mu\text{m}$ |
| Neural network | $7.45 \pm 0.42 \%$ | $0.95 \pm 0.01$ | $5.98 \pm 0.38 \mu\text{m}$ | $9.88 \pm 0.49 \mu\text{m}$ |
| Proposed scaling law | $13.65 \pm 0.68 \%$ | $0.88 \pm 0.02$ | $9.89 \pm 0.62 \mu\text{m}$ | $14.96 \pm 1.14 \mu\text{m}$ |
| Literature scaling law | $17.67 \pm 0.45 \%$ | $0.89 \pm 0.01$ | $10.85 \pm 0.38 \mu\text{m}$ | $14.51 \pm 0.80 \mu\text{m}$ |
| **Generation rate prediction** |  |  |  |  |
| Boosted decision trees | $16.58 \pm 1.05 \%$ | $0.98 \pm 0.00$ | $219.7 \pm 12.7 \text{ Hz}$ | $452.5 \pm 19.5 \text{ Hz}$ |
| Neural network | $22.59 \pm 1.4 \%$ | $0.97 \pm 0.00$ | $260.1 \pm 14.3 \text{ Hz}$ | $499.9 \pm 22.9 \text{ Hz}$ |
| Proposed scaling law | $46.86 \pm 4.60 \%$ | $0.83 \pm 0.03$ | $676.9 \pm 80.7 \text{ Hz}$ | $1182.6 \pm 90.1 \text{ Hz}$ |
| Literature scaling law | $58.68 \pm 3.09 \%$ | $-0.02 \pm 0.24$ | $1367.5 \pm 590.9 \text{ Hz}$ | $2774.7 \pm 582.1 \text{ Hz}$ |

We then trained boosted decision trees to predict normalized droplet diameters, using viscosity ratio, capillary number, flow rate ratio, and the five geometric parameters as inputs. Across 15 randomized training sessions, boosted decision trees showed an MAPE of 5.4% for predicting diameter and 16.6% for generation rate (Fig. 3e, see Supplementary Fig. 14 for 14 additional training sessions). Overall, boosted decision trees (closely followed by the neural network) enabled the most accurate performance prediction in flow-focusing aqueous-in-oil and oil-in-aqueous droplet generation across different fluids with diameters of 15 to 250 *μ*m at rates of 5 to 12000 Hz; models showed higher accuracy for predicting the inner diameter of DEs compared to their outer diameter (Supplementary Fig. 15). Other statistical metrics including coefficient of determination (*R*^2^), mean absolute error (MAE), and root mean square error (RMSE) also demonstrate the significantly higher accuracy of machine learning models compared to scaling laws (Table 3).

Machine learning models show even greater improvements for predicting generation rate. Literature scaling laws resulted in a negative *R*^2^ (i.e., predictions were worse than just predicting the mean outcome for all outcomes) and a MAE of 1367 Hz (MAPE of 58.7%) for predicting generation rate, compared to *R*^2^=0.98 and a MAE of 220 Hz (MAPE of 16.6%) for boosted decision trees and *R*^2^=0.97 and MAE of 260 Hz (MAPE of 22.6%) for neural network.

For both machine learning models, the MAPE for generation rate was approximately 3 times the MAPE for diameter. This is mathematically expected according to conservation of mass. As the generation rate inversely scales with the 3^*rd*^ power of diameter, assuming a relatively small error in diameter prediction and using a Taylor series expansion yields a 3-fold larger MAPE for rate prediction (Supplementary Note 3) [35]. Scaling laws deviate from this rule because their error in predicting diameter is not sufficiently small to neglect higher order approximations in Taylor series expansion.

Boosted decision trees are interpretable and can reveal relative significance of design parameters for a dataset (Methods: Parameter significance study). To determine key parameters in different scenarios of droplet generation, we trained and evaluated decision trees on different subsets of the comprehensive dataset. Flow rate ratio, orifice width, capillary number, and viscosity ratio were most important for predicting normalized droplet diameter in the comprehensive dataset (Fig. 3f.i). For aqueous-in-oil SE droplets, flow rate ratio remained the most important, followed by capillary number (Fig. 3f.ii). For DE droplets, normalized inner diameters were mostly determined by viscosity ratio and flow ratio (Fig. 3f.iii) while normalized outer diameters were affected by all parameters, with capillary number being the most significant (Fig. 3f.iv).

### 2.3 Prediction of stable and unstable DE generation

Producing stable single-core DE droplets requires that generation rates at FF1 and FF2 be matched. If the rate at FF1 exceeds that of FF2, some DEs end up with multiple cores; conversely, if the rate at FF1 is lower than that at FF2, some droplets do not contain a core (Fig. 4a). As generation rates depend critically on device geometries and fluid properties, identifying conditions required to generate stable single-core DEs for new reagent combinations is typically a time-consuming process involving several design iterations and flow rate optimizations for inner, middle, and outer fluids. Here, we tested if machine learning models could streamline this process by predicting device geometries and flow rates required to generate stable single core DEs. The fact that our models can consider DE generation as a combination of 2 independent droplet generation events (i.e. generation of a aqueous-in-oil droplet and an oil-in-aqueous droplet) suggests that these models may also be able to predict when droplet generation is stable (i.e. yields single core DEs) or unstable (i.e. yields multicore or coreless droplets) just by comparing the generation rates at FF1 and FF2.

**Figure 4:**
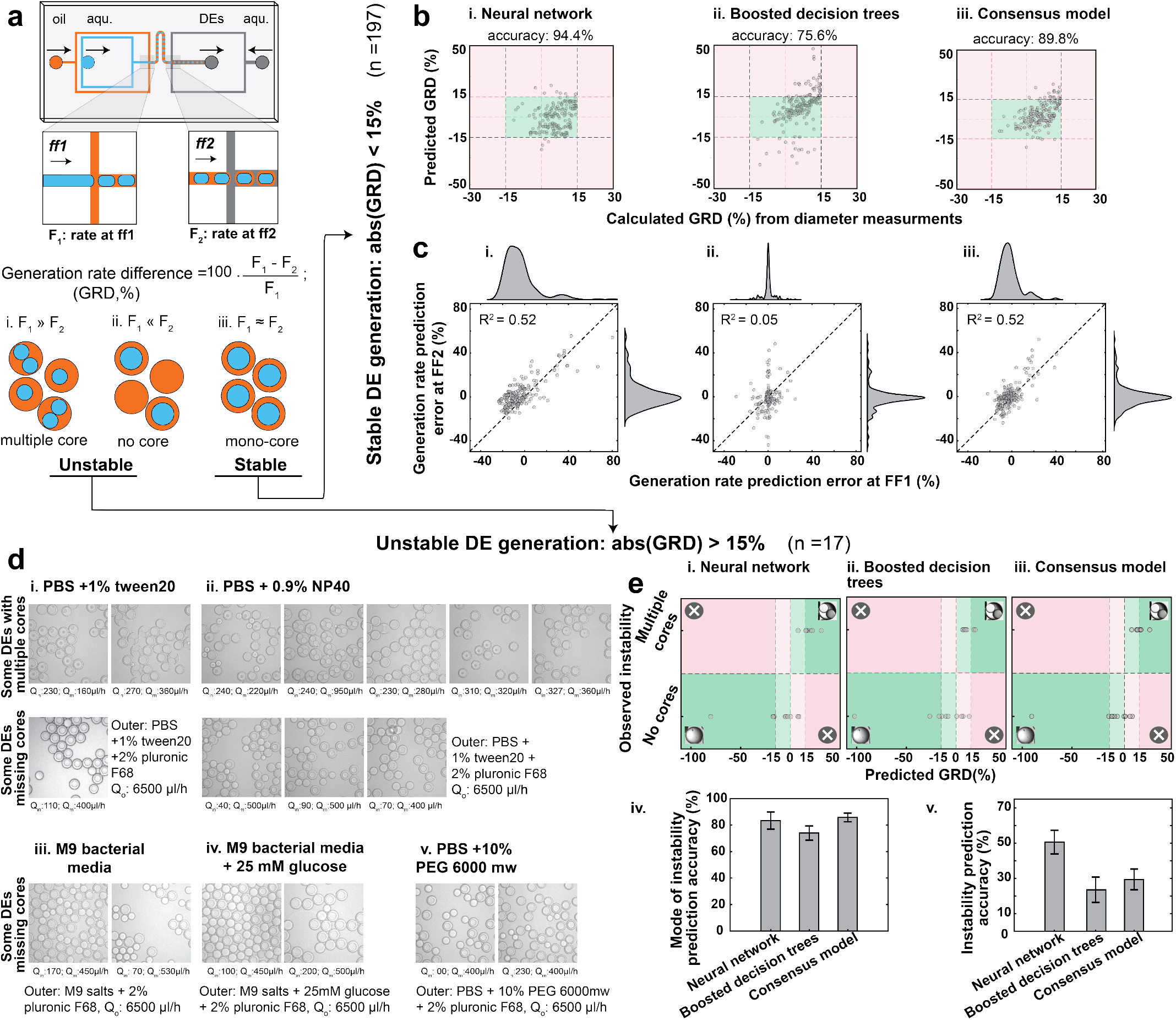
Models’ performance in predicting stable and unstable DE droplet generation regimes. **a**. DE generation was modeled as two events of droplet generation at FF1 (aqueous-in-oil) and FF2 (oil-in-aqueous); the threshold for unstable DE generation was set to a generation rate difference (GRD) of 15%. **b**. Comparison of predicted and calculated GRD at FF1 and FF2 for the neural network, boosted decision trees, and consensus model over the 197 stable datapoints. Green boxes indicate regions with predicted GRD < 15% and experimentally stable droplets; annotated accuracies represent average values over 15 randomized training sessions. **c**. Comparisons between errors in model-predicted generation rates at FF1 and FF2. Markers show comparisons for a single representative model, dashed line indicates 1:1 line, and annotation denotes the average coefficient of determination over 15 randomized training sessions.**d**. Images for unstable DE droplets generated using 17 new conditions (5 different fluid combinations and varied inner and middle flow rates). **e**. Comparisons between observed instability mode vs. predicted GRD (top) and bar charts quantifying accuracy in predicting the mode of instability (bottom, left) and whether or not droplet generation was unstable (bottom, right). Correct predictions appear in green shaded areas, incorrect predictions appear in red shaded areas, and GRDs predicted to lead to stable droplets are indicated by lighter shading. For bar plots, bars indicate average accuracy values across 15 randomized training sessions and error bars indicate standard deviation.

We assessed our machine learning models by (1) predicting the stability of 197 datapoints that resulted in stable DE generation and (2) predicting the instability of 17 newly generated datapoints on unstable DE generation. For the 197 datapoints in the stable DE dataset, we observed a maximum generation rate difference (GRD) of 15% between the experimentally calculated generation rates FF1 and FF2. As the fraction of non-single core DEs depends on the percentage mismatch in generation rates, it is somewhat surprising that mismatches of this magnitude still lead to stable DE generation. This discrepancy likely stems from small inaccuracies in experimentally measured diameters, which scale by a power of 3 when calculating generation rates and the fact that small mismatches still yield droplet populations that are mostly single-core DEs. We therefore classified any set of conditions with a predicted absolute GRD of <15% as yielding stable droplets:

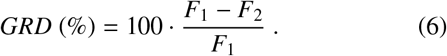

Here, *F*_1_ is generation rate at FF1, calculated from inner diameter, and *F*_2_ is rate at FF2, calculated from outer diameter using conservation of mass.

Using this criteria, the neural network correctly predicted conditions that generate stable, single core DEs for 94.4% of datapoints over 15 randomized training sessions (Fig. 4b.i). Despite predicting diameters and generation rates more accurately than neural networks, boosted decision trees correctly classified conditions as producing stable droplets in only 75.6% of cases (Fig. 4b.ii); in the remaining cases, conditions that generated stable droplets were predicted to be unstable. This performance difference likely stemmed from differing degrees of correlation between model-predicted rate errors at FF1 and FF2 (*R*^2^ = 0.05 and *R*^2^ = 0.52 for boosted decision trees and neural networks, respectively, Fig. 4c.i–ii). To take advantage of the high accuracy of boosted decision trees in predicting generation rates and the high accuracy of the neural network in predicting DE stability, we developed a consensus model that averages predictions of each model. This consensus model correctly predicted stability for 89.8% of data-points while also minimizing generation rate errors (Fig. 4b.iii & c.iii).

Next, we tested if these models could predict unstable DE generation despite only being trained on conditions that lead to stable DE generation within the comprehensive dataset. We generated 17 new datapoints using 5 different fluids to make DE droplets with either multiple or no cores (Fig. 4d). We then tested if these models correctly predicted unstable generation (i.e. if they predicted a GRD >15%). While machine learning models fairly accurately predicted the mode of instability (i.e., GRD >0 : multiple cores or GRD <0 : no cores) for unstable droplets (neural networks, boosted decision trees, and the consensus model classified modes of instability correctly in 83.5%, 74.1%, and 85.8% of cases, respectively, Fig. 4e), they were less able to predict whether or not DE generation was stable (neural networks, boosted decision trees, and the consensus model predicted absolute *GRD* > 15% for 50.6%, 23.5%, and 29.4% of unstable generation cases, respectively) (Fig. 4e). This prediction performance could likely be improved in the future by training on a dataset including a much larger number of unstable generation cases.

### 2.4 Machine learning models can generalize to previously unseen geometries and fluids

Training models that accurately generalize to unseen design parameters and new data sources is a common challenge in developing machine learning models [45–47]. Here, we directly tested the ability of each model (the literature scaling model, the neural network, the boosted decision tree, and the consensus model) to generalize by using each model to predict droplet diameter, generation rate, and stability for as-yet-unseen fluids and device geometries by (1) comparing model predictions of droplet diameter and generation rate for previously published data not included in the training data and (2) using each model to predict diameter and generation rate and then experimentally fabricating devices with specified new geometries, using them to generate droplets using those fluids, and directly comparing model predictions with new experimental results. This evaluation of the accuracy of ‘blind’ model predictions provides a stringent test of the degree to which each model can generalize.

First, we used two previously published datasets with unseen geometries and fluids to evaluate the generalizability of our models to datasets generated by others (Fig. 5a.i). These datasets include SE and DE aqueous-in-oil [24] and oil-in-polymer droplets [48]. In the aqueous-in-oil SE data, droplets of DMEM mammalian cell media with added 16% optiprep, 10% FBS, and 1% penicillin-streptomycin were generated with HFE 7500 oil containing 1.5% fluorinated surfactant for single-cell analysis [24]. In the oil-in-polymer data [48], coreshell structures were formed using HFE 7500, trimethylolpropane trimethacrylate (TMPTMA), and 50% glycerol in DI water as inner, middle, and outer fluids, respectively. As fluid properties and interfacial tension were only provided for FF1 (droplets of HFE 7500 in TMPTMA) in this dataset, we predicted only inner diameter of DEs. Predicted droplet diameters for both datasets (18 datapoints total) were least accurate for the literature scaling law (MAPE of 21.7% and 45.0% for diameter and rate, respectively). The machine learning models were consistently more accurate, with boosted decision trees slightly outperforming others in terms of MAPE (10.6% for diameter and 28.6% for rate) and the consensus model slightly outperforming others in terms of coefficient of determination (*R*^2^ of 0.97 and 0.98 for diameter and rate, respectively) averaged over 15 randomized training sessions (Fig. 5a.ii–iv).

**Figure 5:**
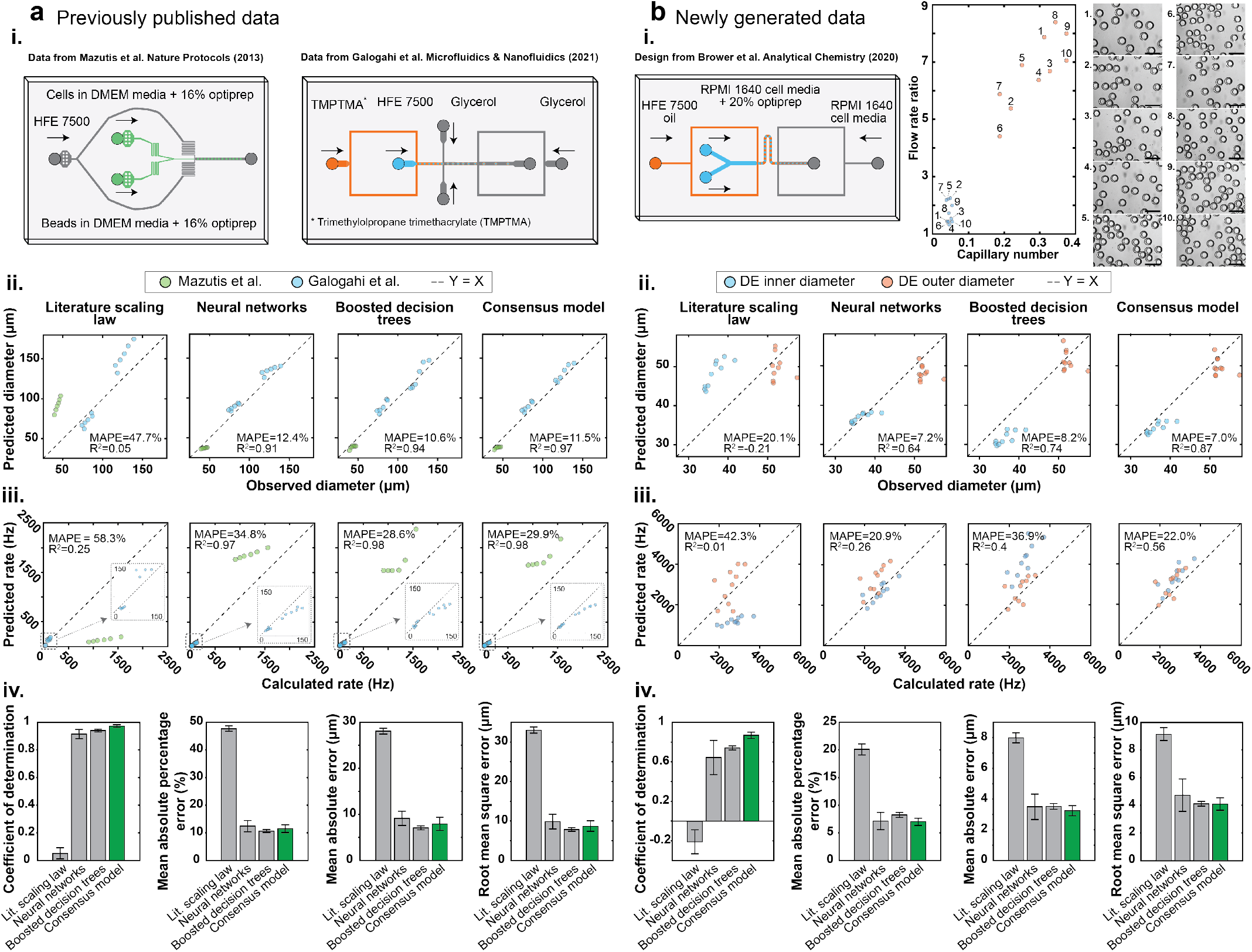
Generalization of machine learning models to fluids and geometries not included within the comprehensive dataset. **a.i** Performance of trained models in predicting previously published SE droplet geometries and generation rates not included within the training set. **a.i**References for experimental data and schematics showing device geometries and fluid compositions used to generate SE droplets. **a.ii-iii** Comparison between observed SE droplet diameters (a.ii) and generation rates (a.iii) and those predicted by 4 different trained models. Dashed lines indicate 1:1 identity line. **a.iv** Quality metrics assessing model performance. Bars indicate average performance across 15 randomized training sessions; error bars denote standard deviation. **b** Performance of trained model in predicting DE geometries and generation rates for newly generated data. **b.i**. Schematics showing device geometry, fluid compositions, flow rate ratios, and capillary numbers used to generate DEs (left) and representative images of generated DEs (right). Scale bars are 50 *μ*m. **b.ii-iii** Comparison between observed DE droplet diameters (b.ii) and generation rates (b.iii) and those predicted by 4 different trained models. Dashed lines indicate 1:1 identity line. **b.iv** Quality metrics assessing model performance. Bars indicate average performance across 15 randomized training sessions; error bars denote standard deviation.

Next, we fabricated a new DE generation device based on a previously published design [20] with as-yet-unseen channel geometries (2 aqueous inlets instead of a single inlet and a normalized channel depth of 1.33 instead of 1) and used it to generate DEs with fluids suitable for mammalian cell encapsulation (complete RPMI 1640 cell media with 20% optiprep and 0.1% pluronic F-127 for inner fluid, and complete RPMI 1640 with 5% pluronic F-127 for the outer fluid). After measuring the interfacial surface tension for each fluid interface (required to calculate capillary number), we used each of the pre-trained models to predict droplet diameters and generation rates for 10 different flow rate combinations. Finally, we generated DEs using these same flow rates and directly compared model predictions to experimental data. The resultant droplets spanned inner diameters of 29.6 to 34.9 *μ*m and outer diameters of 36.7 to 46.8 *μ*m (Fig. 5b.i). The machine learning models all outperformed the literature scaling law and accurately predicted droplet diameters and generation rates (MAPEs of 7.0 to 8.2% and 20.9 to 36.9% for diameter and generation rate, respectively), with the consensus model showing the best overall performance (Fig. 5b.ii–iv). This ability to accurately predict newly generated data with unseen fluids and geometries demonstrates an ability to generalize, likely due to the diversity of the comprehensive dataset in terms of geometry, fluid properties, and flow rates and the use of dimensionless parameters and *L*2 regularization during model training [49].

### 2.5 Design automation of SE and DE droplets

The ability to automate design of devices for producing droplets with desired diameters and generation rates can dramatically reduce time spent fabricating, testing, and optimizing microfluidic devices. We previously developed an online open-source tool (DAFD, for **D**esign **A**utomation of **F**luid **D**ynamics) that converted user-specified droplet diameters and rates into a microfluidic design and flow rates that delivered the desired performance [35]. However, the previous version of this tool was limited to only aqueous-in-oil SEs, a single simple fluid combination (DI water and mineral oil), large polycarbonate devices (orifice width > 75 *μ*m), and a maximum generation rate of 500 Hz. Here, we present a new open-source-tool, DAFD 3.0, that leverages the consensus model (i.e., average of neural network and boosted decision trees) and new search algorithms to design PDMS and polycarbonate devices capable of producing aqueous-in-oil and oil-in-aqueous SE and DE droplets using a wide variety of different fluids. This tool supports device orifice widths of 15 to 175 *μ*m and droplets of 15 to 250 *μ*m in diameter produced at rates of 5 to 12000 Hz.

For SE design automation, DAFD 3.0 takes the desired diameter and rate alongside the viscosities and interfacial tension of inner and outer fluids as inputs and provides the necessary device geometry and flow rates (while allowing for optional design constraints; Supplementary Fig. 16 and Methods). To test DAFD 3.0 accuracy and reliability for SEs, we specified that we wanted to produce SEs with diameters of 25, 30, and 35 *μ*m using a previously unseen fluid combination (RPMI 1640 complete cell media with added 20% optiprep and 0.1% pluronic F127 as dispersed fluid and dSurf HFE 7500 as the continuous fluid) and constrained the possible geometry to require the same pre-fabricated DE generator device used to assess model generalizability in (Fig. 5.b). We then generated SEs with the DE generator by blocking the outer fluid inlet and flowing dispersed and continuous fluids through FF1 and compared resulting droplet diameters to our original specifications. Introducing fluids using DAFD-suggested flow rates (Supplementary Table 4) yielded SEs of 27.5, 31.6, and 37.9 *μ*m in diameter, very close to model predictions with an overall MAE of 2.36 *μ*m and MAPE of 7.94% (Fig. 6.a).

**Figure 6:**
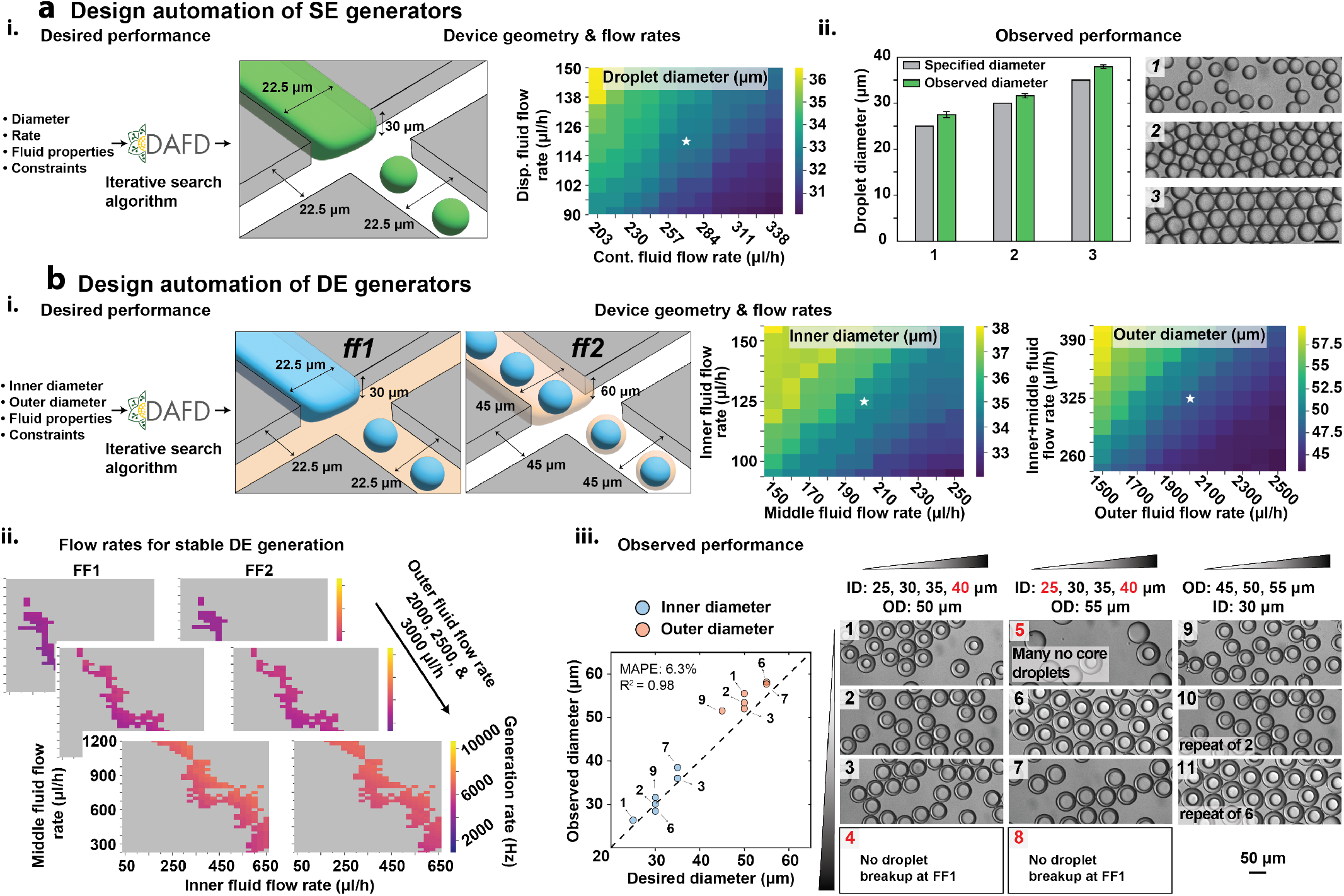
Trained machine learning models and custom search algorithms enable design automation of SE and DE droplet generation. **a** Design automation of SE droplet generation. **a.i** DAFD 3.0 takes user-specified diameter, rate, fluid properties, and optional constraints as inputs and returns the necessary geometry and flow rates required to generate the desired droplets. **a.ii**. DAFD-predicted and measured droplet diameters after specifying desired SE diameters of 25, 30, and 35 *μ*m for an unseen fluid combination (left) and representative images of generated droplets (right). Measured droplets differed from specified droplets by a MAE of 2.3 *μ*m (MAPE of 7.9%). **b** Design automation of DE droplet generation. **b.i**. DAFD 3.0 also converts user-specified DE inner and outer diameters to the necessary geometries and flow rates required to generate them. **b.ii**. DAFD-predicted generation rates as a function of middle, inner, and outer flow rates; predict generation rate differences (GRDs) between FF1 and FF2 to identify likely stable (GRD <5%) and unstable (GRD >5%) regimes. **b.iii** Comparison between observed and DAFD-specified DE inner (blue) and outer (orange) diameters for an unseen fluid combination and 9 different flow rates (left); images show representative DE droplets generated under each condition (right). For stable droplets, observed inner and outer diameters differed from those specified by a MAE of 2.7 *μ*m (MAPE of 6.3%). Scale bars are 50 *μ*m.

For DE design automation, our tool takes desired inner and outer diameters and fluid properties (viscosities and interfacial tensions) of 3 fluids as inputs and predicts DE inner and outer diameters generated using either six different default designs or a user-specified geometry (if a suitable solution can be found). For DE generation, DAFD 3.0 requires the total flow rate at FF1 (i.e., the inner plus middle fluid flow rates) to be equal to the flow rate of dispersed fluid at FF2 to uphold conservation of mass. Next, generation rates at FF1 and FF2 are calculated by the tool, and only the datapoints that have a GRD of less than 5% are considered to increase the chances of stable DE generation and establish a solution space (Fig. 6b.ii). Finally, DAFD 3.0 ranks potential solutions based on their average deviation from the desired inner and outer diameters and returns a single best set of flow rates (Supplementary Fig. 17 and Methods).

To test DAFD 3.0 accuracy and reliability for DEs, we specified: (1) that we wanted to generate DEs with a range of inner (25–40 *μ*m) and outer diameters (45–55 *μ*m) using the same fluids as used above for SEs within an outer fluid of 5% pluronic F127 and (2) that we wanted to constrain the design to the same DE device geometry used above. We then used the 9 suggested flow rate combinations to generate DEs and quantified the resultant droplet diameters (Supplementary Table 5). Consistent with prior observations that machine learning models can struggle to accurately predict whether flow combinations yield stable single-core droplets, 3/9 conditions at the extremes of inner droplet diameter did not yield stable droplets. The flow rates designed to create DEs with the largest inner diameter (40 *μ*m ID and either 50 or 55 *μ*m OD) led to no droplet formation at FF1, while the flow rates designed to create DEs with a 25 *μ*m ID and a 55 *μ*m OD led to many DEs lacking a core. Among stable datapoints, DAFD 3.0 was highly accurate, generating DEs that were different from target diameters by a MAE of 2.70 *μ*m (MAPE of 6.3%). The accuracy for inner diameter (MAE of 1.5 *μ*m & MAPE of 4.8%) was higher than outer diameter (MAE of 3.9 *μ*m & MAPE of 7.9%), potentially due to the bias of training data toward aqueous-in-oil droplets and the non-zero yet minimal dependence of outer diameter on the inner diameter (Fig. 6b.iii).

## 3 Discussion

Here, we establish that machine learning models can enable accurate and generalizable prediction of droplet diameters and generation rates based on device geometries, fluid properties, and flow rates for aqueous-in-oil and oil-in-aqueous SE and DE droplets. These models cover droplet diameters of 15 to 250 *μ*m at rates of up to 12000 Hz and take a broad range of input design parameters, including 3 orders of magnitude variation in capillary number, 2 orders of magnitude variation in viscosity ratio, and more than 1 order of magnitude variation in flow rate ratio and microfluidic channel size. This represents a notable improvement over previous models for single-fluid aqueous-in-oil SEs that could either only account for variations in geometry for generation rates of up to 500 Hz [35] or surfactants for a single geometry [36]. In all cases, the trained neural network and boosted decision trees detailed here outperform previously published scaling laws and machine learning models in terms of accuracy and parameter range [35, 36, 38–43]. Boosted decision trees had the highest accuracy, with a MAE of 4.6 *μ*m (MAPE of 5.4%) and 220 Hz (16.6%) in predicting diameter and generation rate, respectively. Nonetheless, a consensus model based on both boosted decision trees and neural networks resulted in better generalizability to as-yet-unseen fluids and geometries.

Our models account for variations in fluid properties, flow rates, and geometry by taking 7 dimensionless inputs (capillary number, flow rate ratio, viscosity ratio, normalized channel depth, normalized dispersed fluid inlet width, normalized continuous fluid inlet width, and normalized outlet width) and orifice width as the only input with units. The dimensionless inputs and a dimensionless output (droplet diameter normalized by hydraulic diameter of orifice) enable generalizability to newly created and previously published data that the models were not trained on. Our models’ accuracy in predicting stable and unstable DE generation demonstrates their ability to cover both aqueous-in-oil and oil-in-aqueous droplets and validates our simplifying assumption that DE generation can be modeled as two independent events of droplet generation with a non-zero yet minimal loss in accuracy.

Predictive models can be integrated with custom search algorithms to create design automation tools that translate user-specified SE and DE characteristics and input fluids to the necessary geometry and flow rates. Our tool also enables rapid performance characterization of droplet generators. For instance, for a given DE generator all flow rates that result in stable DE generation can be quickly mapped to find its deliverable range of inner and outer diameters and generation rates. Using the consensus model that averages the predictions of neural networks and boosted decision trees, we packaged performance prediction and design automation as an online and open-source software tool to eliminate the need for design iterations when developing SE and DE generators. DAFD 3.0 requires the viscosities and interfacial tension of fluids used for droplet generation, which can be readily measured using standard rheological and in-situ techniques [30, 50] or approximated using the properties of similar fluids.

The generalizable predictive power of our models is partly due to leveraging two independently created datasets for microfluidic droplet generation. We therefore envision future versions of DAFD to benefit from new publicly available datasets to achieve higher accuracy and account for broader range of parameters. As a result future repositories for microfluidic data in addition to repositories for device designs such as Metafluidics [51] would greatly benefit community-driven design automation efforts. Future integration of our tool with other computer-aided design tools for microfluidics [52, 53] and real-time dynamic control schemes [54, 55] would enable automated conversion of high-level user-specifications to fabrication-ready designs that robustly deliver the desired performance.

Sophisticated high-throughput microfluidic operations require multiple integrated components to function optimally in tandem. As the number of on-chip components increases, the possible design space grows exponentially, making designing and optimizing such platforms extremely challenging [31]. A predictive understanding of microfluidic components can be leveraged to achieve new functionalities more complex than that of its individual components. We establish that a predictive understanding of SE generators can be leveraged to achieve performance prediction and design automation of DE generators (a two-component device). Similarly, we expect that a predictive understanding of droplet generators in conjunction with other components such as deterministic lateral displacement arrays, inertial focusers, pico-injectors, and cell and droplet sorters to enable novel high-throughput screening platforms that are developed in a matter of days instead of several months. Our work paves the way for custom and highly optimized microfluidic platforms to be readily adapted to new applications to accelerate the discovery process in life science.

## 4 Methods

### 4.1 Dimensionless numbers and flow rate calculations

Droplet generation can be considered as a competition between viscous forces exerted by the continuous fluid and cohesive forces within the dispersed fluid [28]. As a result, the capillary number (given by the ratio of viscous forces to interfacial tension) is commonly used to describe the characteristics of droplet generation. In flow-focusing droplet generation, capillary number can be defined as given in Eq. (7):

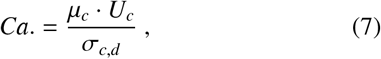

where μ_*c*_ is the dynamic viscosity of the continuous fluid, *U*_*c*_ is the characteristic velocity of the continuous fluid through the orifice (flow rate of the continuous fluid divided by the cross-sectional area of orifice), and σ_*c,d*_ is the interfacial tension between the continuous and dispersed fluids.

The flow rate of the continuous fluid can therefore be calculated using capillary number, fluid properties, and device geometry as given in Eq. (8):

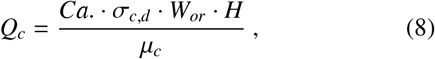

where *Q*_*c*_ is the flow rate of the continuous fluid, *W*_*or*_ is the orifice width, and *H* is channel depth.

The flow rate of the dispersed fluid can be determined from the flow rate of the continuous fluid and the flow rate ratio as follows:

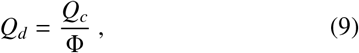

where *Q*_*d*_ is the dispersed fluid flow rate, *Q*_*c*_ is the flow rate of the continuous fluid, and Φ is the flow rate ratio.

Some of the predictive models developed here also take viscosity ratio as an input, defined using Eq. (10):

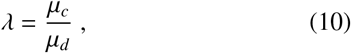

where μ_*c*_ is the dynamic viscosity of the continuous fluid and μ_*d*_ is the dynamic viscosity of the dispersed fluid.

To improve the generalizability of our models to different size scales, we converted the geometric parameters of a flow-focusing device to dimensionless numbers by normalizing them by the orifice width (except for orifice width itself);

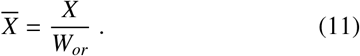

Here, 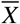 is the dimensionless (normalized) geometric parameter, and *X* can be channel height (*H*), dispersed inlet width (*DIW*), continuous inlet width (*CIW*), or outlet channel width (*OCW*). Orifice length was not considered as a design parameter in our models due to its negligible effect on droplet size and rate in the dripping regime. This also enables us to model flow-focusing geometries with unclear orifice lengths (e.g., when the outlet channel has the same width as the orifice width such that the orifice length could equally be considered to be 0 or equal to the length of outlet channel).

### 4.2 Measurement of fluid properties

Standard pendant drop tensiometry with drop shape analysis was used to measure the interfacial tension between different pairs of dispersed and continuous fluids, as reported in Table 1 and as previously described [30, 56]. Since the density of HFE 7500 oil is greater than the density of inner and outer fluids used in this study, pendant oil droplets were suspended within the inner or outer aqueous fluids to measure the interfacial tension. A metal capillary nozzle (27 gauge) was used to suspend oil droplets within 5 mL of inner or outer fluid. A custom MATLAB code was used to analyze the oil droplet shape and calculate interfacial tension, as previously established [57, 58]. Briefly, shape analysis was conducted when the oil droplet was as stable as possible. Since droplets were observed at equilibrium, cohesive forces (interfacial tension) and gravitational deformation are balanced and the simplified Young-Laplace equation can be equated to hydrostatic pressure and solved to estimate interfacial tension. All reported interfacial tension measurements were the average of 3–6 analyzed drops. The dynamic viscosity of fluids was also measured using a commercial rotational cone and plate rheometer, as previously described [30]. Briefly, a 2 ° cone at 20° C was used to conduct a logarithmic flow sweep across a broad range of shear rates (2.86479–2864.79 Hz). The average viscosity in the linear regime was reported as the shear rate-independent viscosity of the fluid.

### 4.3 Device fabrication

SE droplet generators were rapidly prototyped using a low-cost desktop micromill (Bantam Tools) to ablate microfluidic channels with the smallest dimension of 75 *μ*m out of a polycarbonate substrate (McMaster-Carr), as previously described [59]. Once channel geometries were etched into polycarbonate, plastic burrs and dirt were removed by using a soft brush followed by sonication in IPA and DI water. Devices were then sealed using a thin layer of PDMS (Sylgard 184) sandwiched between two polycarbonate layers or through an 81 *μ*m double-sided adhesive (ArCare 90445, Adhesive Research) and then placed in a vacuum desiccator to remove air bubbles and complete device sealing.

DE droplet generators were fabricated through standard photolithography followed by soft-lithography as previously described [30]. Briefly, a silicon wafer with two different heights (the height at flow-focuser 2 is double the height at flow-focuser 1) was created using 2-layer SU8 deposition and standard photo-lithography [20]. To cast PDMS devices from this master mold, we poured a 1:5 ratio of PDMS on the wafer, degassed, and cured for 15 minutes at 80°c. Inlets and out-lets were punched using a 1 mm biopsy punch (Robbins Instruments) and then this featured layer was placed on a blank slab of 1:10 PDMS (that was cured for 15 minutes at 80°C) and baked for 48 hours at 80°C to bond the device to the blank slab (via off-ratio bonding) and render the PDMS device hydrophobic (longer bake times result in smaller pore sizes within PDMS and improve its hydrophobicity).

### 4.4 Single emulsion generation

Single emulsions were generated using a microfluidic device made out of polycarbonate using a low-cost desktop micromill, as previously described [35]. DI water with added food coloring for better visualization was used as the dispersed fluid. NF 350 mineral oil with a viscosity of 57.2 mPa.s and density of 857 kg/m^3^ was used as the continuous fluid and 5% V/V Span-80 surfactant (Sigma-Aldrich) was added to the oil to stabilize droplets. Fluids were introduced into the microfluidic device using syringe pumps (Harvard Apparatus) and syringes were connected to the device through PVC tubing (McMaster-Carr). All fluids were filtered using 0.45 *μ*m polyvinylidene fluoride (PVDF) membrane filters (Millipore) before loading. Once flow rates were set on the syringe pumps, we waited for 10 minutes before collecting droplets to ensure flow stability. Oil was always introduced before DI water to ensure that the continuous fluid (i.e., oil) wet the surfaces of the channels first. Droplets were imaged using a high-speed camera (IDT Xstream) mounted on a stereo-microscope (AmScope) and experiments were illuminated using an 18,000 Lumen LED light source (Expert Digital Imaging) placed underneath the microfluidic device. Once a high-speed video of an experiment was recorded, we analyzed it using an open-source custom python code we previously developed to record droplet diameter, generation rate, and droplet polydispersity: https://github.com/CIDARLAB/uDrop-Generation. The dataset on SE generation is available in an Open Science Framework repository: https://osf.io/938rs/.

### 4.5 Double emulsion generation

Double emulsions were generated using PDMS microfluidic devices fabricated as described above [30]. A variety of inner and outer fluids commonly used in life science applications were used as inner and outer fluids. HFE 7500 fluorinated oil (Sigma-Aldrich) with a viscosity of 1.6 mPa.s with added 2.2% ionic PEG-Krytox surfactant (FSH 157, Miller-Stephenson) was used as the middle fluid (i.e., oil). The surfactant added to the outer fluid varied depending on its properties (as detailed in Table 1); in most experiments, we added 2% Pluronic F-68 with or without added 1% Tween-20, except for the complete RPMI 1640 cell media experiments, where only 5% added Pluronic F-127 (Sigma-Aldrich) was used to stabilize the DEs. The majority of DE datapoints used to create a comprehensive dataset and initially train models were taken from our previous study using a single device geometry [30].

We also generated new data for droplets with a broader range of inner and outer DE diameters to test model prediction accuracy. These new data were generated using 2 additional DE generation devices that were either scaled down or scaled up versions of the original device (orifice width at FF1 set to 15 or 30 *μ*m (instead of 22.5 *μ*m) and orifice width at FF2 set to 30 or 60 *μ*m (instead of 45 *μ*m) while keeping normalized channel depth, normalized outlet width, and normalized dispersed and continues fluid inlet widths to 1). Prior to running experiments, each device was surface treated to render the 2^*nd*^ half of the device (FF2) hydrophilic. This was achieved by taping (Scotch tape) over the inner and middle fluids inlets (to protect the 1^*st*^ half of the device, FF1, from being exposed to plasma) and allowing air/oxygen plasma to enter through the outer fluid inlet and the device outlet, thereby rendering the FF2 region of the device hydrophilic (10 minutes of plasma treatment)[20, 60]. Fluids were introduced to the microfluidic device using syringe pumps (Harvard Apparatus) using 0.015” I.D. and 0.043” O.D. LDPE polyethylene medical tubing (BB31695-PE/2, Scientific Commodities). All fluids were filtered using 0.45 *μ*m polyvinylidene fluoride (PVDF) membrane filters (Millipore) before loading. Immediately after surface treatment, we flowed the outer fluid (sheath) into the devices to ensure that the flow-focuser 2 region of the device remained hydrophilic; 30 seconds after the introduction of the outer fluid, we introduced the middle fluid (oil) and inner fluid. The flow rates of the middle and inner fluids were initialized with a value higher than the intended final value to speed fluid entry into the flow-focusers and then slowly lowered to the intended flow rates. Once flow rates were set on syringe pumps, we waited 4-minute intervals before collecting droplets to ensure flow stability. Droplet generation was imaged using a high-speed camera (ASI174MM, ZWO) mounted on a stereo-microscope (AmScope). Once DEs were collected, they were imaged inside a cell-counter chamber slide (Countess) using an inverted microscope and a custom MATLAB-based image processing workflow to measure the inner and outer diameters of the DEs, as previously described [30]. The image processing source code is available at: https://osf.io/pt6qu/?view_only=f1690e6efd7a4773b7e26fec5a65aada. The dataset on DE generation is available as an Open Science Framework repository: https://osf.io/938rs/.

### 4.6 Training machine learning models

#### Neural network

The comprehensive dataset used to train models is relatively limited (∼1000 datapoints) compared to datasets traditionally used to train deep neural networks. We therefore used a shallow neural network comprised of two hidden layers of 512 and 16 nodes with rectified linear units (ReLU) activation functions [44]. We trained the model to minimize a mean squared error loss using an Adam optimizer with a learning rate of 0.0003 and batches of size 32 [61]. We also L2 regularized the model parameters with a penalty term of 0.001 to prevent the model from overfitting [49]. This model took 7 design parameters as inputs (orifice width plus 6 dimensionless numbers: capillary number, flow rate ratio, normalized channel depth, normalized dispersed fluid inlet width, normalized continuous fluid inlet width, and normalized outlet channel width). The model then predicted a dimensionless droplet diameter (normalized by the hydraulic diameter of the orifice) as the output. We did not include viscosity ratio as an input parameter for the neural network since it resulted in slightly lower accuracy in predicting unseen previously published data, despite resulting in slightly higher accuracy when predicting the comprehensive dataset.

#### Boosted decision trees

We used the XGBoost package for implementing the boosted decision trees [62]. Our model consists of 100 boosted trees, trained to minimize a mean squared error loss with an L2 regularization penalty term of 1 and a learning rate of 0.3. To prevent individual trees from overfitting, we limited tree depth to 6 and halted leaf node splitting once their weight was below 1. This model took 8 design parameters as inputs (orifice width plus 7 dimensionless numbers: viscosity ratio, capillary number, flow rate ratio, normalized channel depth, normalized dispersed fluid inlet width, normalized continuous fluid inlet width, and normalized outlet channel width). The model then predicted a dimensionless droplet diameter (normalized by the hydraulic diameter of the orifice) as the output.

#### Evaluation

We assessed models by randomly partitioning the comprehensive dataset into train and test sets, comprising 80% and 20% of the original dataset, respectively. Table 3 displays the average accuracy metrics against the test set across 15 randomized training sessions. The scatter plots of predictions for all figures are shown based a single model predictions that had a prediction performance close to the average performance of 15 randomized training sessions. The source codes for training, testing, and validating the neural network and the boosted decision trees are available on Open Science Framework: https://osf.io/938rs/ and our GitHub repository: https://github.com/CIDARLAB/DAFD-website.

### 4.7 Parameter significance study

For boosted decision trees, we defined parameter significance as the average loss reduction across the node splits where the parameter serves as the decision variable (also referred to as gain in XGBoost). We calculated each parameters’ significance for 15 randomized training sessions and reported their averaged significance in Fig. 4f.i. Moreover, we repeated our evaluation of parameter significance on subsets of the comprehensive dataset, constraining each subset to only include certain datapoint types: single emulsions, double-emulsion inner diameters, and double-emulsion outer diameters, as depicted in Figures 4f. ii., iii., and iv., respectively.

### 4.8 Single emulsion design automation

In DAFD 3.0, users first select whether to generate a design for single or double emulsions. For single emulsions, the user enters the desired droplet diameter and/or generation rate, rheological properties of fluids (e.g., viscosities, interfacial tension), and any constraints to the geometric design or flow conditions of the droplet generator. A custom iterative optimization algorithm is then used to find a design and flow rates that deliver the desired performance, as described previously [35]. First, the closest experimental data point is found that fits the design constraints. If this closest point fits all constraints and produces a diameter and rate within 3 *μ*m and 15*Hz*, respectively, the experimental point is returned without design iterations. If the closest point is not within these ranges, an iterative optimization process is implemented. For a maximum of 5000 iterations, each parameter is stepped up or down by a specific amount, unless this parameter is constrained or this new value passes the preset parameter bounds (Supplementary Table 6). In this workflow, the average prediction of both the neural network and boosted decision trees are used to predict droplet diameter. Prediction accuracy is determined by a model error cost function:

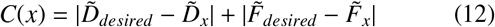

where *C*(*x*) is the cost of a design *x*, 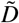 is the scalar normalized droplet diameter, and 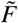 is the scalar normalized generation rate. Both droplet diameter and generation rate are normalized to a standard scalar to prevent bias from the larger range of possible generation rates (hundreds to thousands) compared to diameters (tens to few hundred). If the user only specifies diameter or rate, only that value is included in the cost function. This process is repeated until the cost function reaches zero, the maximum number of iterations is reached, or the change in the cost function is less than a preset tolerance of 10^−9^. The geometric design parameters and the flow conditions of the final solution is then returned to the user alongside the predicted droplet diameter and generation rate. Additionally, the predicted diameters and generation rates for flow rates up to ±25% of the designed value are provided to construct a performance heat map as a device-specific operation guideline for users. The source code for our SE design automation algorithm is available on our GitHub repository: https://github.com/CIDARLAB/DAFD-website.

### 4.9 Double emulsion design automation

A separate approach is implemented for design automation of double emulsion generators (since this requires pairing two droplet generators in series). First, the user provides the desired inner and outer diameters and the rheological properties of their desired inner, middle, and outer fluids. Next, six preset devices can be selected, which have orifice widths of 15 *μ*m, 22.5 *μ*m, and 30 *μ*m at FF1 (i.e., orifice widths of 30, 45, 60 *μ*m at FF2, respectively) and a normalized channel depth of 1 or 1.33. If none of the six devices are selected by the user, all are considered in the design automation workflow. The user can also specify a custom double emulsion generator geometry if preferred. After taking the user inputs, the entire flow space of the two droplet generators is simulated (50–650 μL/hr for the inner aqueous fluid, 200–1200 μL/hr for the middle fluid, and 1500–10000 μL/hr for the outer aqueous fluid). The dispersed flow rate of flow-focuser 2 (FF2) is simulated across all unique combinations of the total flow rate of FF1 (inner aqueous fluid flow rate plus the middle fluid flow rate) to ensure that the final design is compatible with conservation of mass. Each of the data points of FF1 is then paired with points from FF2 that have matching flow conditions and less than a 5% predicted generation rate difference (GRD). Any designs outside of a 5% GRD are deemed unstable and excluded from consideration. No solution is returned if no points with a GRD less than 5% are found. The pairings with GRD < 5% are then ranked according to the total percentage error in their predicted inner and outer diameters from the user-specified values. Top candidate designs are then recommended to the user in case a certain generation rate is preferred or an error in the inner or outer diameter is more tolerable. The source code for DAFD 3.0 and the design automation workflow is available at https://github.com/CIDARLAB/dafd-website.

### 4.10 Data availability

The comprehensive dataset and its subsets (SE and DE datasets) are available as an Open Science Framework: https://osf.io/938rs/ and DAFD’s website at: http://dafdcad.org.

### 4.11 Code availability

All source code generated and used in this study for performance prediction and design automation SE and DE droplets are available at: https://github.com/CIDARLAB/DAFD-website

## Acknowledgements

A.L. acknowledges funding as a Damon Runyon Postdoctoral Fellow (DRG-2479-22) from Damon Runyon Cancer Research Foundation. D.P.M. acknowledges funding from the Society of Lab Automation and Screening Graduate Education Fellowship. S.G.K.C. acknowledges funding as a ChEM-H CBI fellow and a Stanford Graduate SGF fellow. D.M.D. and D.P.M. are supported by NSF Semiconductor Synthetic Biology for Information Storage and Retrieval (Award #2027045). P.M.F. is a Chan Zuckerberg Biohub investigator and this work was partly supported by NIH DP2 GM123641 awarded to P.M.F., a Stanford Bio-X interdisciplinary seed grant, and the Emerson Collective.

## Author contributions

A.L. established the research idea, created the SE dataset, trained models, designed and carried out experiments, and analyzed data. D.P.M. helped establish the research idea, developed the algorithms for design automation, and incorporated performance prediction and design automation for DAFD website. S.G.K.C. measured all fluid properties and created the DE dataset. K.K. helped with training machine learning models and improved their generalizability. D.M.D. helped establish the research idea, supported the generation of the SE dataset and development of DAFD, and partially funded the project. P.M.F. helped establish the research idea, supported the generation of the DE dataset, and oversaw and funded the project. All authors read and approved this manuscript.

## Competing interests

The authors declare no competing interests.

## Additional information

### Supplementary Information

Information on range of capillary numbers and flow rate ratios included in this study is provided in Supplementary Fig. 1. Information on previously published experimental datasets and machine learning approaches for flow-focusing droplet size prediction is provided in Supplementary Table 1. Additional information on previously published scaling laws, their performance against the comprehensive data, and their constants when fitted to our dataset can be found in Supplementary Note 1, Supplementary Table 2, and Supplementary Fig. 2–12. Droplet diameter and generation rate prediction accuracies for 14 consecutive training sessions for the neural network and boosted decision trees can be found in Supplementary Fig 13 & 14, respectively. Information on including viscosity ratio as a design parameter of the neural network is provided in Supplementary Note 2 and Supplementary Table 3. A comparison of the accuracies of our machine learning models in predicting inner and outer diameters of DEs in our dataset is provided in Supplementary Figure 15. Schematic overview of the algorithms developed for design automation of SE and DE generators are provided in Supplementary Figures 16 & 17, respectively. The input desired performance and constraints and the DAFD suggested geometry and flow rates for SE and DE design automation examples are provided in Supplementary Tables 4 & 5, respectively. The parameter range and step-size for SE design automation is provided in Supplementary Table 6.

## Notes

https://osf.io/938rs/

### Competing Interest Statement

The authors have declared no competing interest.

## References

[1] Lauren D Zarzar, Vishnu Sresht, Ellen M Sletten, Julia A Kalow, Daniel Blankschtein, and Timothy M Swager. Dynamically reconfigurable complex emulsions via tunable interfacial tensions. Nature, 518(7540):520–524, 2015.

[2] Iain C Clark, Michael A Wheeler, Hong-Gyun Lee, Zhaorong Li, Liliana M Sanmarco, Shravan Thaploo, Carolina M Polonio, Seung Won Shin, Giulia Scalisi, Amy R Henry, et al. Identification of astrocyte regulators by nucleic acid cytometry. Nature, pages 1–3, 2023.

[3] Grace XY Zheng, Jessica M Terry, Phillip Belgrader, Paul Ryvkin, Zachary W Bent, Ryan Wilson, Solongo B Ziraldo, Tobias D Wheeler, Geoff P McDermott, Junjie Zhu, et al. Massively parallel digital transcriptional profiling of single cells. Nature communications, 8(1):1–12, 2017.

[4] Rapolas Zilionis, Juozas Nainys, Adrian Veres, Virginia Savova, David Zemmour, Allon M Klein, and Linas Mazutis. Single-cell barcoding and sequencing using droplet microfluidics. Nature protocols, 12(1):44–73, 2017.

[5] Fabrice Gielen, Raphaelle Hours, Stephane Emond, Martin Fischlechner, Ursula Schell, and Florian Hollfelder. Ultrahigh-throughput– directed enzyme evolution by absorbance-activated droplet sorting (aads). Proceedings of the National Academy of Sciences, 113(47):E7383–E7389, 2016.

[6] Derek Vallejo, Ali Nikoomanzar, Brian M Paegel, and John C Chaput. Fluorescence-activated droplet sorting for single-cell directed evolution. ACS synthetic biology, 8(6):1430–1440, 2019.

[7] Katherine S Elvira, Robert CR Wootton, Andrew J deMello, et al. The past, present and potential for microfluidic reactor technology in chemical synthesis. Nature chemistry, 5(11):905–915, 2013.

[8] Gisbert Schneider. Automating drug discovery. Nature reviews drug discovery, 17(2):97–113, 2018.

[9] Annabelle Gérard, Adam Woolfe, Guillaume Mottet, Marcel Reichen, Carlos Castrillon, Vera Menrath, Sami Ellouze, Adeline Poitou, Raphaël Doineau, Luis Briseno-Roa, et al. High-throughput single-cell activity-based screening and sequencing of antibodies using droplet microfluidics. Nature biotechnology, 38(6):715–721, 2020.

[10] Andrew S Utada, Elise Lorenceau, Darren R Link, Peter D Kaplan, Howard A Stone, and DA Weitz. Monodisperse double emulsions generated from a microcapillary device. Science, 308(5721):537–541, 2005.

[11] Bárbara Herranz-Blanco, Laura R Arriaga, Ermei Mäkilä, Alexandra Correia, Neha Shrestha, Sabiruddin Mirza, David A Weitz, Jarno Salonen, Jouni Hirvonen, and Hélder A Santos. Microfluidic assembly of multistage porous silicon–lipid vesicles for controlled drug release. Lab on a Chip, 14(6):1083–1086, 2014.

[12] Jenni Pessi, Hélder A Santos, Inna Miroshnyk, David A Weitz, Sabiruddin Mirza, et al. Microfluidics-assisted engineering of polymeric micro-capsules with high encapsulation efficiency for protein drug delivery. International journal of pharmaceutics, 472(1-2):82–87, 2014.

[13] Justin E Silpe, Janine K Nunes, Albert T Poortinga, and Howard A Stone. Generation of antibubbles from core–shell double emulsion templates produced by microfluidics. Langmuir, 29(28):8782–8787, 2013.

[14] Yves Hennequin, Nicolas Pannacci, Concepción Pulido De Torres, Georgios Tetradis-Meris, Stephane Chapuliot, Elisabeth Bouchaud, and Patrick Tabeling. Synthesizing microcapsules with controlled geometrical and mechanical properties with microfluidic double emulsion technology. Langmuir, 25(14):7857–7861, 2009.

[15] Milla G Santos, Débora A Carpinteiro, Marcelo Thomazini, Glaucia A Rocha-Selmi, Adriano G da Cruz, Christiane EC Rodrigues, and Carmen S Favaro-Trindade. Coencapsulation of xylitol and menthol by double emulsion followed by complex coacervation and microcapsule application in chewing gum. Food research international, 66:454–462, 2014.

[16] Margot Stasse, Tiphaine Ribaut, Véronique Schmitt, and Valérie Héroguez. Encapsulation of lipophilic fragrance by polymerization of the intermediate aqueous phase of an oil-in-water-in-oil (o/w/o) double emulsion. Polymer Chemistry, 10(30):4154–4162, 2019.

[17] Anastasia Zinchenko, Sean RA Devenish, Balint Kintses, Pierre-Yves Colin, Martin Fischlechner, and Florian Hollfelder. One in a million: flow cytometric sorting of single cell-lysate assays in monodisperse picolitre double emulsion droplets for directed evolution. Analytical chemistry, 86(5):2526–2533, 2014.

[18] Evan Z Macosko, Anindita Basu, Rahul Satija, James Nemesh, Karthik Shekhar, Melissa Goldman, Itay Tirosh, Allison R Bialas, Nolan Kamitaki, Emily M Martersteck, et al. Highly parallel genome-wide expression profiling of individual cells using nanoliter droplets. Cell, 161(5):1202–1214, 2015.

[19] Yun Ding, Philip D Howes, and Andrew J deMello. Recent advances in droplet microfluidics. Analytical chemistry, 92(1):132–149, 2019.

[20] Kara K Brower, Margarita Khariton, Peter H Suzuki, Chris Still, Gaeun Kim, Suzanne GK Calhoun, Lei S Qi, Bo Wang, and Polly M Fordyce. Double emulsion picoreactors for high-throughput single-cell encapsulation and phenotyping via facs. Analytical chemistry, 92(19):13262–13270, 2020.

[21] Wei Wang, Mao-Jie Zhang, and Liang-Yin Chu. Microfluidic approach for encapsulation via double emulsions. Current opinion in pharmacology, 18:35–41, 2014.

[22] Alphonsus HC Ng, Songming Peng, Alexander M Xu, Won Jun Noh, Katherine Guo, Michael T Bethune, William Chour, Jongchan Choi, Sung Yang, David Baltimore, et al. Mate-seq: microfluidic antigen-tcr engagement sequencing. Lab on a Chip, 19(18):3011–3021, 2019.

[23] Shelley L Anna, Nathalie Bontoux, and Howard A Stone. Formation of dispersions using “flow focusing” in microchannels. Applied physics letters, 82(3):364–366, 2003.

[24] Linas Mazutis, John Gilbert, W Lloyd Ung, David A Weitz, Andrew D Griffiths, and John A Heyman. Single-cell analysis and sorting using droplet-based microfluidics. Nature protocols, 8(5):870–891, 2013.

[25] AR Abate, A Poitzsch, Y Hwang, J Lee, J Czerwinska, and DA Weitz. Impact of inlet channel geometry on microfluidic drop formation. Physical Review E, 80(2):026310, 2009.

[26] Shengqing Xu, Zhihong Nie, Minseok Seo, Patrick Lewis, Eugenia Kumacheva, Howard A Stone, Piotr Garstecki, Douglas B Weibel, Irina Gitlin, and George M Whitesides. Generation of monodisperse particles by using microfluidics: control over size, shape, and composition. Angewandte Chemie, 117(5):734–738, 2005.

[27] Shelley Lynn Anna. Droplets and bubbles in microfluidic devices. Annual Review of Fluid Mechanics, 48:285–309, 2016.

[28] Charles N Baroud, Francois Gallaire, and Rémi Dangla. Dynamics of microfluidic droplets. Lab on a Chip, 10(16):2032–2045, 2010.

[29] Stefan Wiedemeier, Marko Eichler, Robert Römer, Andreas Grodrian, Karen Lemke, Krees Nagel, Claus-Peter Klages, and Gunter Gastrock. Parametric studies on droplet generation reproducibility for applications with biological relevant fluids. Engineering in life sciences, 17(12):1271–1280, 2017.

[30] Suzanne GK Calhoun, Kara K Brower, Vineeth Chandran Suja, Gaeun Kim, Ningning Wang, Alexandra L McCully, Halim Kusumaatmaja, Gerald G Fuller, and Polly M Fordyce. Systematic characterization of effect of flow rates and buffer compositions on double emulsion droplet volumes and stability. Lab on a Chip, 22(12):2315–2330, 2022.

[31] Elishai Ezra Tsur. Computer-aided design of microfluidic circuits. Annu. Rev. Biomed. Eng, 22:285–307, 2020.

[32] David McIntyre, Ali Lashkaripour, Polly Fordyce, and Douglas Densmore. Machine learning for microfluidic design and control. Lab on a Chip, 22(16):2925–2937, 2022.

[33] Adam R Abate, Julian Thiele, and David A Weitz. One-step formation of multiple emulsions in microfluidics. Lab on a Chip, 11(2):253–258, 2011.

[34] Sangam Srikanth, Satish Kumar Dubey, Arshad Javed, and Sanket Goel. Droplet based microfluidics integrated with machine learning. Sensors and Actuators A: Physical, 332:113096, 2021.

[35] Ali Lashkaripour, Christopher Rodriguez, Noushin Mehdipour, Rizki Mardian, David McIntyre, Luis Ortiz, Joshua Campbell, and Douglas Densmore. Machine learning enables design automation of microfluidic flow-focusing droplet generation. Nature communications, 12(1):1–14, 2021.

[36] Loïc Chagot, César Quilodrán-Casas, Maria Kalli, Nina M Kovalchuk, Mark JH Simmons, Omar K Matar, Rossella Arcucci, and Panagiota Angeli. Surfactant-laden droplet size prediction in a flow-focusing microchannel: a data-driven approach. Lab on a Chip, 22(20):3848–3859, 2022.

[37] Ali Lashkaripour, Christopher Rodriguez, Luis Ortiz, and Douglas Densmore. Performance tuning of microfluidic flow-focusing droplet generators. Lab on a Chip, 19(6):1041–1053, 2019.

[38] Zheyu Liu, Maojie Chai, Xin Chen, Seyed Hossein Hejazi, and Yiqiang Li. Emulsification in a microfluidic flow-focusing device: Effect of the dispersed phase viscosity. Fuel, 283:119229, 2021.

[39] Taotao Fu, Yining Wu, Youguang Ma, and Huai Z Li. Droplet formation and breakup dynamics in microfluidic flow-focusing devices: From dripping to jetting. Chemical engineering science, 84:207–217, 2012.

[40] Haihu Liu and Yonghao Zhang. Droplet formation in microfluidic cross-junctions. Physics of Fluids, 23(8):082101, 2011.

[41] Thomas Ward, Magalie Faivre, Manouk Abkarian, and Howard A Stone. Microfluidic flow focusing: Drop size and scaling in pressure versus flow-rate-driven pumping. Electrophoresis, 26(19):3716–3724, 2005.

[42] Wingki Lee, Lynn M Walker, and Shelley L Anna. Role of geometry and fluid properties in droplet and thread formation processes in planar flow focusing. Physics of Fluids, 21(3):032103, 2009.

[43] Jian Hong Xu, SW Li, Jing Tan, and GS Luo. Correlations of droplet formation in t-junction microfluidic devices: from squeezing to dripping. Microfluidics and Nanofluidics, 5:711–717, 2008.

[44] Xia Hu, Lingyang Chu, Jian Pei, Weiqing Liu, and Jiang Bian. Model complexity of deep learning: A survey. Knowledge and Information Systems, 63:2585–2619, 2021.

[45] Kenji Kawaguchi, Leslie Pack Kaelbling, and Yoshua Bengio. Generalization in deep learning. arXiv preprint arXiv:1710.05468, 2017.

[46] Behnam Neyshabur, Srinadh Bhojanapalli, David McAllester, and Nati Srebro. Exploring generalization in deep learning. Advances in neural information processing systems, 30, 2017.

[47] Joachim Krois, Anselmo Garcia Cantu, Akhilanand Chaurasia, Ranjitkumar Patil, Prabhat Kumar Chaudhari, Robert Gaudin, Sascha Gehrung, and Falk Schwendicke. Generalizability of deep learning models for dental image analysis. Scientific reports, 11(1):1–7, 2021.

[48] Fariba Malekpour Galogahi, Yong Zhu, Hongjie An, and Nam-Trung Nguyen. Formation of core–shell droplets for the encapsulation of liquid contents. Microfluidics and Nanofluidics, 25(10):1–11, 2021.

[49] Corinna Cortes, Mehryar Mohri, and Afshin Rostamizadeh. L2 regularization for learning kernels. arXiv preprint arXiv:1205.2653, 2012.

[50] Rosa D’Apolito, Antonio Perazzo, Mariapia D’Antuono, Valentina Preziosi, Giovanna Tomaiuolo, Reinhard Miller, and Stefano Guido. Measuring interfacial tension of emulsions in situ by microfluidics. Langmuir, 34(17):4991–4997, 2018.

[51] David S Kong, Todd A Thorsen, Jonathan Babb, Scott T Wick, Jeremy J Gam, Ron Weiss, and Peter A Carr. Open-source, community-driven microfluidics with metafluidics. Nature biotechnology, 35(6):523–529, 2017.

[52] Radhakrishna Sanka, Joshua Lippai, Dinithi Samarasekera, Sarah Nem-sick, and Douglas Densmore. 3d μ f-interactive design environment for continuous flow microfluidic devices. Scientific reports, 9(1):9166, 2019.

[53] Haiyao Huang and Douglas Densmore. Fluigi: microfluidic device synthesis for synthetic biology. ACM Journal on Emerging Technologies in Computing Systems (JETC), 11(3):1–19, 2014.

[54] Alexander E Siemenn, Evyatar Shaulsky, Matthew Beveridge, Tonio Buonassisi, Sara M Hashmi, and Iddo Drori. A machine learning and computer vision approach to rapidly optimize multiscale droplet generation. ACS Applied Materials & Interfaces, 14(3):4668–4679, 2022.

[55] Oliver J Dressler, Philip D Howes, Jaebum Choo, and Andrew J deMello. Reinforcement learning for dynamic microfluidic control. ACS omega, 3(8):10084–10091, 2018.

[56] Nicolas J Alvarez, Lynn M Walker, and Shelley L Anna. A non-gradient based algorithm for the determination of surface tension from a pendant drop: Application to low bond number drop shapes. Journal of colloid and interface science, 333(2):557–562, 2009.

[57] V Chandran Suja, Mariana Rodriguez-Hakim, Javier Tajuelo, and Gerald G Fuller. Single bubble and drop techniques for characterizing foams and emulsions. Advances in Colloid and Interface Science, 286:102295, 2020.

[58] Jianghu Cui, Yunhua Yang, Yonghui Hu, and Fangbai Li. Rice husk based porous carbon loaded with silver nanoparticles by a simple and cost-effective approach and their antibacterial activity. Journal of colloid and interface science, 455:117–124, 2015.

[59] Ali Lashkaripour, Ryan Silva, and Douglas Densmore. Desktop micromilled microfluidics. Microfluidics and Nanofluidics, 22(3):1–13, 2018.

[60] Kara K Brower, Catherine Carswell-Crumpton, Sandy Klemm, Bianca Cruz, Gaeun Kim, Suzanne GK Calhoun, Lisa Nichols, and Polly M Fordyce. Double emulsion flow cytometry with high-throughput single droplet isolation and nucleic acid recovery. Lab on a Chip, 20(12):2062–2074, 2020.

[61] Zijun Zhang. Improved adam optimizer for deep neural networks. In 2018 IEEE/ACM 26th International Symposium on Quality of Service (IWQoS), pages 1–2. Ieee, 2018.

[62] Tianqi Chen and Carlos Guestrin. Xgboost: A scalable tree boosting system. In Proceedings of the 22nd acm sigkdd international conference on knowledge discovery and data mining, pages 785–794, 2016.

