## Supplementary information for "Design Automation of Microfluidic Single and Double Emulsion Droplets with Machine Learning"

May 31, 2023

#### Supplementary Material

Supplementary Table 1: Comparison of previously published experimental datasets and machine learning models for predicting droplet diameter in microfluidic flow-focusing droplet generation in the dripping regime.

| **Parameter** | Chagot et al. (2022) [1] | Lashkaripour et al. (2021) [3] | This study |
|---|---|---|---|
| Number of datapoints | 468 | 474 | 868 |
| Range of diameters | 73 - 190 $\mu$m | 25 - 250 $\mu$m | 15 - 250 $\mu$m |
| Range of generation rates | 5 - 800 Hz | 5 - 500 Hz | 5 - 12000 Hz |
| Variations in geometry | No | Yes | Yes |
| Variations in fluids | No | No | Yes |
| Variations in surfactants | Yes | No | Yes |
| Aqueous-in-oil droplets | Yes | Yes | Yes |
| Oil-in-aqueous droplets | No | No | Yes |
| Single/Double emulsions | Single | Single | Single & Double |
| Dimensionless inputs | No | Yes | Yes |
| Dimensionless output | Yes | No | Yes |
| Tool for performance prediction | No | Yes | Yes |
| Tool for design automation | No | Yes | Yes |

Supplementary Table 2: Fitting constants for all the scaling laws tested against our comprehensive dataset on SE and DE droplet generation.

| Scaling law | Fitting constants | | | | | |
|---|---|---|---|---|---|---|
| | $c_1$ | $c_2$ | $c_3$ | $c_4$ | $c_5$ | $c_6$ |
| $\frac{D}{D_{hyd}} = c_1 Ca^{c_2}$[4] | 0.590 | -0.043 | N/A | N/A | N/A | N/A |
| $\frac{D}{W_{or}} = c_1(\frac{Q_d}{Q_c})^{c_2}$[7] | 1.579 | 0.373 | N/A | N/A | N/A | N/A |
| $\frac{D}{W_{or}} = c_1(\frac{Q_d}{Q_c})^{c_2} Ca_c^{c_3}$ [2] | 1.566 | 0.347 | -0.062 | N/A | N/A | N/A |
| $\frac{D}{D_{hyd}} = c_1(\frac{Q_d}{Q_c})^{c_2} Ca_c^{c_3}$ | 1.534 | 0.373 | -0.009 | N/A | N/A | N/A |
| $\frac{D}{W_c} = (c_1 + C_2 \frac{Q_d}{Q_c}) Ca^{c_3}$ [5] | 0.162 | 0.447 | -0.080 | N/A | N/A | N/A |
| $\frac{D}{W_{or}} = (c_1 + C_2 \frac{Q_d}{Q_c}) Ca^{c_3}$ | 0.538 | 1.507 | -0.047 | N/A | N/A | N/A |
| $\frac{D}{W_{or}} = c_1 + c_2(\frac{Q_d}{Q_c})^{c_3} Ca^{c_4}$ [8] | -6.687 | 8.183 | 0.043 | -0.0066 | N/A | N/A |
| $\frac{D}{D_{hyd}} = c_1 + c_2(\frac{Q_d}{Q_c})^{c_3} Ca^{c_4}$ | -0.303 | 1.784 | 0.273 | -0.012 | N/A | N/A |
| $\frac{D}{D_{hyd}} = c_1(\frac{Q_d}{Q_c})^{c_2}(\frac{\lambda_d}{\lambda_c})^{c_3} Ca_c^{c_4}$ [6] | 1.768 | 0.333 | -0.105 | -0.089 | N/A | N/A |
| $\frac{D}{D_{hyd}} = c_1 + c_2(\frac{Q_d}{Q_c})^{c_3}(c_4 + Ca^{c_5})(\frac{\lambda_c}{\lambda_d})^{c_6}$ | -0.173 | 0.010 | 0.293 | 189.93 | 1.313 | -0.040 |

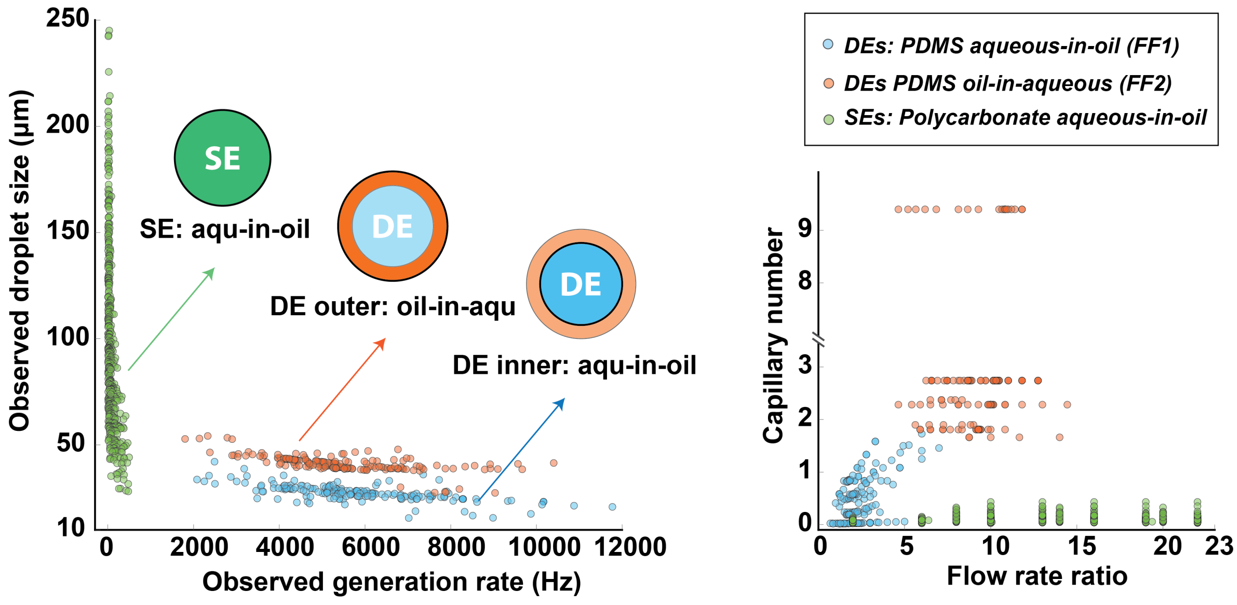


Supplementary Figure 1: Distribution of capillary numbers and flow rate ratios in the comprehensive dataset.

### 1 Note 1: Literature scaling laws

To establish the baseline for performance prediction accuracy, we fitted a number of scaling laws from the literature to our comprehensive dataset and assessed their accuracy in predicting droplet diameter and generation rate.

First, we considered the Fu et al. scaling law [2], where a dimensionless droplet size (i.e., droplet size normalized by the oil inlet width) is proposed to scale with flow rate ratio and capillary number, as given in Eq. (1):

$$\frac{L}{W_{or}} = c_1(\frac{Q_d}{Q_c})^{c_2} Ca_c^{c_3} \tag{1}$$

Here, $L$ is droplet size, $W_{or}$ is the orifice width, $Q_d$ is the flow rate of the dispersed fluid, $Q_c$ is the flow rate of the continuous fluid, and $Ca_c$ is the capillary number of the continuous fluid. This model showed a MAPE of 21.7% in predicting the droplet diameter and 88.9% in predicting generation rate of the 20% test set in the comprehensive dataset.

We next attempted to improve the Fu et al. scaling law by normalizing the droplet diameter with the hydraulic diameter instead of the orifice width, as given in Eq. (2).

$$\frac{L}{D_{hyd}} = c_1(\frac{Q_d}{Q_c})^{c_2} Ca_c^{c_3} \tag{2}$$

This improved model predicted droplet diameter with a MAPE of 17.2% and generation rate with a MAPE of 78.6% for the test set of the comprehensive dataset, with an example

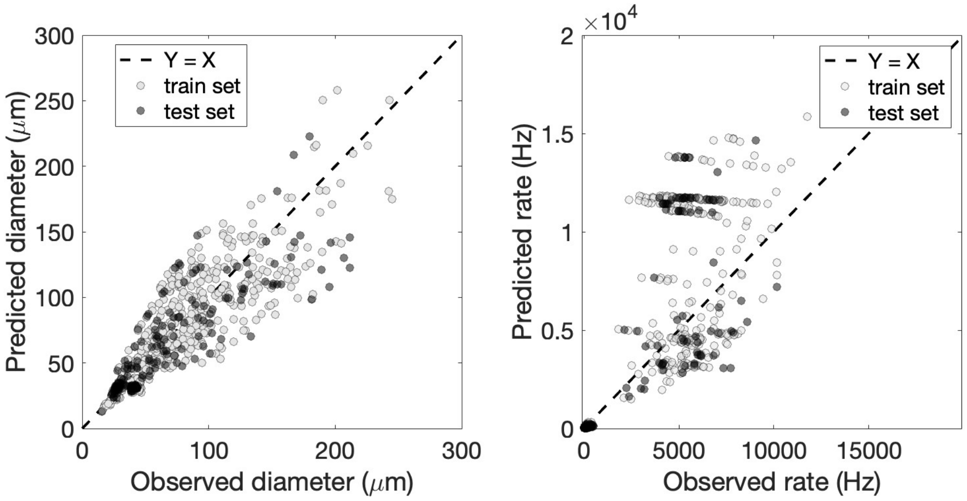


Supplementary Figure 2: Fu at el. scaling law.

run shown in Fig. S.3.

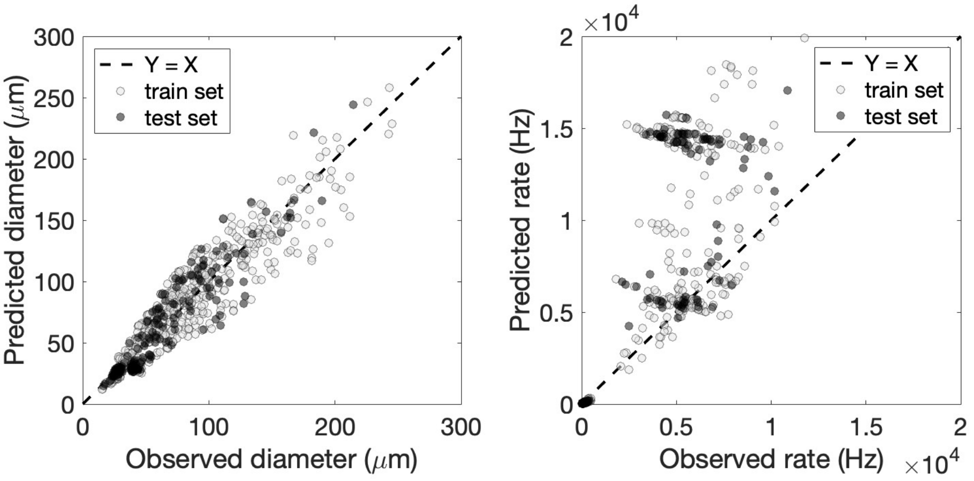


Supplementary Figure 3: Improved scaling law adapted from Fu et al. scaling law.

Next, we considered the Liu et al. scaling law [5], where droplet diameter normalized by the oil inlet width scales with a capillary number and flow rate ratio as given in Eq. (4):

$$\frac{D}{W_c} = (c_1 + C_2 \frac{Q_d}{Q_c}) Ca^{c_3} \tag{3}$$

This scaling law was not accurate and resulted in a MAPE of 47.6% in predicting droplet size and a MAPE of 3023% in predicting the generation rates of the test set, as shown in Fig. S.(4).

Additionally, we considered an improved version of the Liu et al. scaling law [5] in which the droplet diameter is normalized by orifice width instead of the oil inlet width, as given in

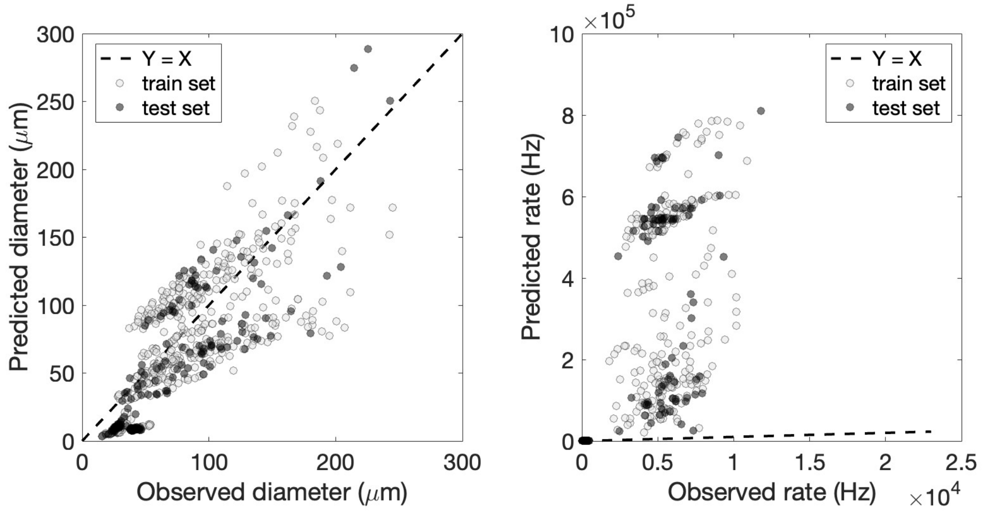


Supplementary Figure 4: Scaling law proposed by Liu et al.

Eq. (4).

$$\frac{D}{W_{or}} = (c_1 + C_2\frac{Q_d}{Q_c})Ca^{c_3} \tag{4}$$

The improved version of the Liu et al. scaling law showed a MAPE of 24.4% and 100.7% in predicting the droplet diameter and generation rate of the comprehensive test set, as shown in Fig. S5.

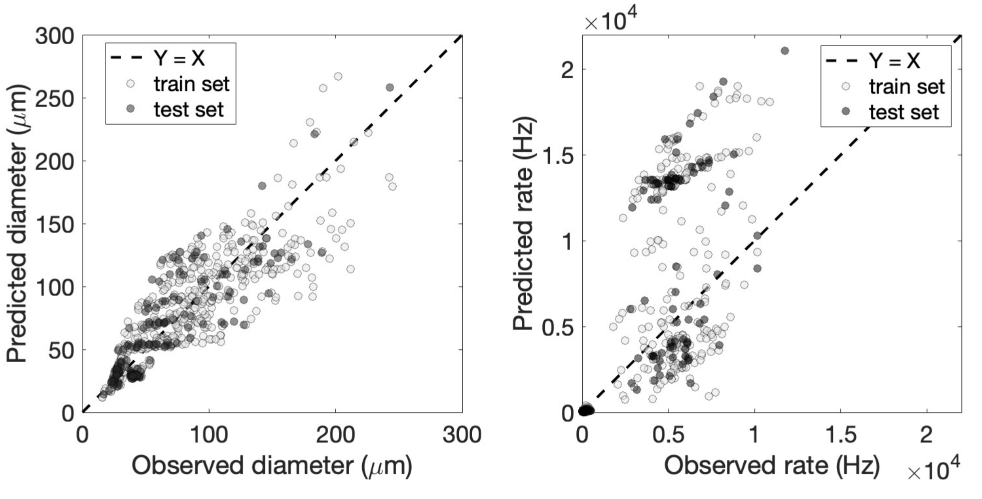


Supplementary Figure 5: An improved version of the scaling law originally proposed by Liu et al.

Next, we fitted a scaling law proposed by Xu et al. [8], where the normalized droplet diameter scales non-linearly with flow rate ratio and capillary number, as given in Eq. (5).

$$\frac{D}{W_{or}} = c_1 + c_2(\frac{Q_d}{Q_c})^{c_3}Ca^{c_4} \tag{5}$$

This scaling law predicted the droplet diameters and generation rates in the test set of the comprehensive dataset with a MAPE of 20.7% and 83.5%, respectively. An example test is shown in Fig. S. 6.

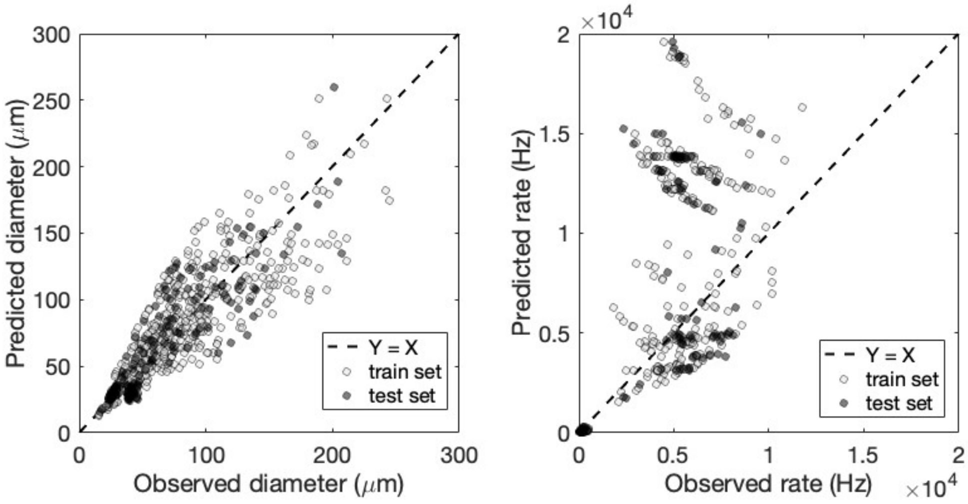


Supplementary Figure 6: Scaling law originally proposed by Xu et al.

We then improved the scaling law proposed by Xu et al. [8] by normalizing the droplet diameter by the hydraulic diameter instead of the orifice width, as given in Eq. (6).

$$\frac{D}{D_{hyd}} = c_1 + c_2(\frac{Q_d}{Q_c})^{c_3} Ca^{c_4} \tag{6}$$

The improved scaling law was capable of predicting droplet diameter with a MAPE of 17.2% and generation rate with a MAPE of 78.5% within the test set of the comprehensive dataset, as shown in Fig. S.7.

Next, we considered a scaling law proposed by Lee et al. [4], where the droplet diameter normalized by the hydraulic diameter scales with capillary number, as given in Eq. 7.

$$\frac{D}{D_{hyd}} = c_1 Ca^{c_2} \tag{7}$$

This scaling law predicted the droplet diameters and generation rates in the test set of the comprehensive dataset with a MAPE OF 34.9% and 324%, respectively, as shown in Fig. S.8.

Next, we fitted a scaling law proposed by Ward et al. [7], where the droplet diameter normalized by the orifice scales with the ratio of dispersed and continuous flow rates, as given in Eq. (8).

$$\frac{D}{W_{or}} = c_1(\frac{Q_d}{Q_c})^{c_2} \tag{8}$$

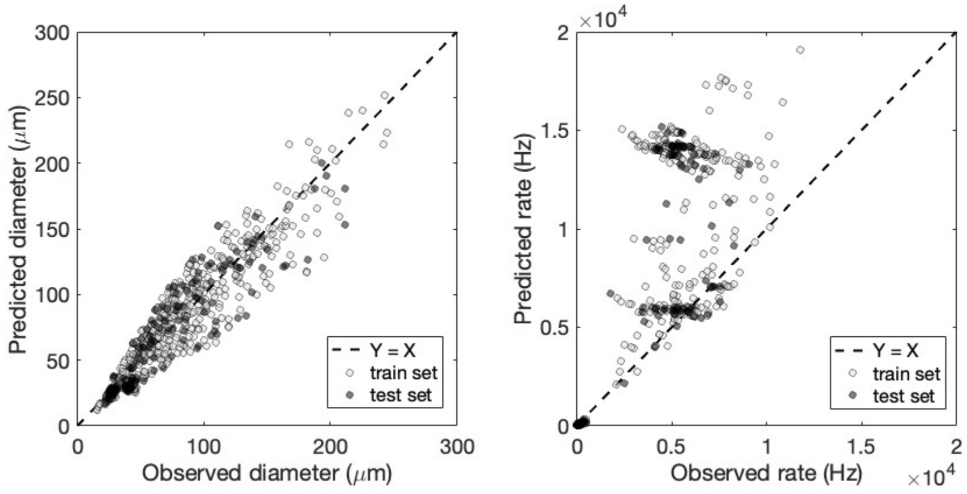


Supplementary Figure 7: An improved version of the scaling law originally proposed by Xu et al.

This scaling law predicted the droplet diameter with a 21.3% MAPE and generation rate with a 139% MAPE for the datapoints in the test set of the comprehensive dataset, as shown in Fig. S.9.

Next, we fitted a scaling law proposed by Zheya Liu et al. [6] to our data. In this scaling law the normalized droplet diameter by the hydraulic diameter scales with capillary number, flow rate ratio, and viscosity ratio, as given in Eq. (9).

$$\frac{D}{D_{hyd}} = c_1(\frac{Q_d}{Q_c})^{c_2}(\frac{\lambda_d}{\lambda_c})^{c_3}Ca_c^{c_4} \tag{9}$$

This scaling law demonstrated the highest accuracy of all previously published scaling laws in predicting our comprehensive dataset, with a MAPE of 17.7% in predicting the diameter and 58.7% in predicting the generation rate, as shown in Fig. S.10.

Finally, we proposed a new scaling law that takes capillary number, flow rate ratio, and viscosity ratio to predict the droplet diameter normalized by the hydraulic diameter of the orifice, as given in Eq. 10.

$$\frac{D}{D_{hyd}} = c_1 + c_2(\frac{Q_d}{Q_c})^{c_3}(c_4 + Ca^{c_5})(\frac{\lambda_c}{\lambda_d})^{c_6} \tag{10}$$

The proposed scaling law was able to predict the droplet diameter with a MAPE of 13.6% and generation rate with a MAPE of 46.9%, as shown in Fig. S.11.

A single comparison of the MAPE in predicting droplet size and rate for all the scaling laws is depicted in Fig. S.12.

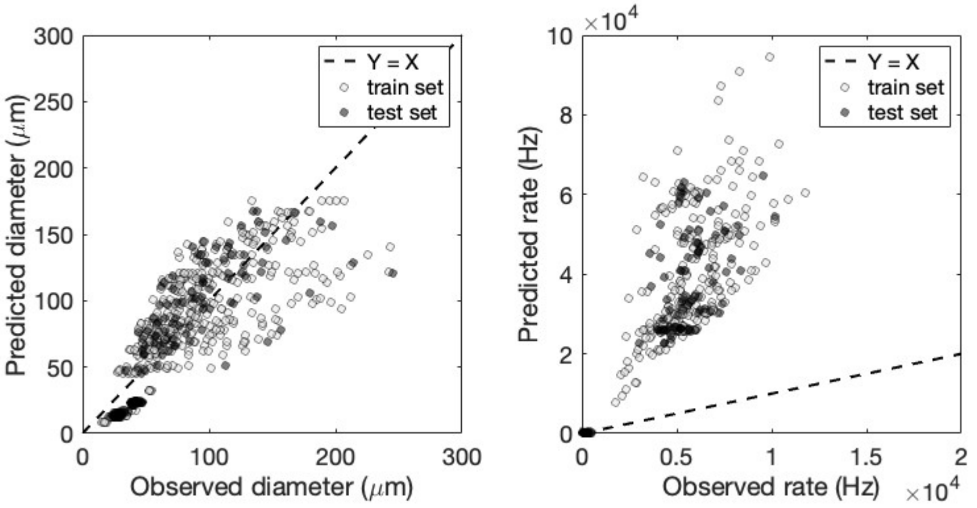


Supplementary Figure 8: The scaling law proposed by Lee et al.

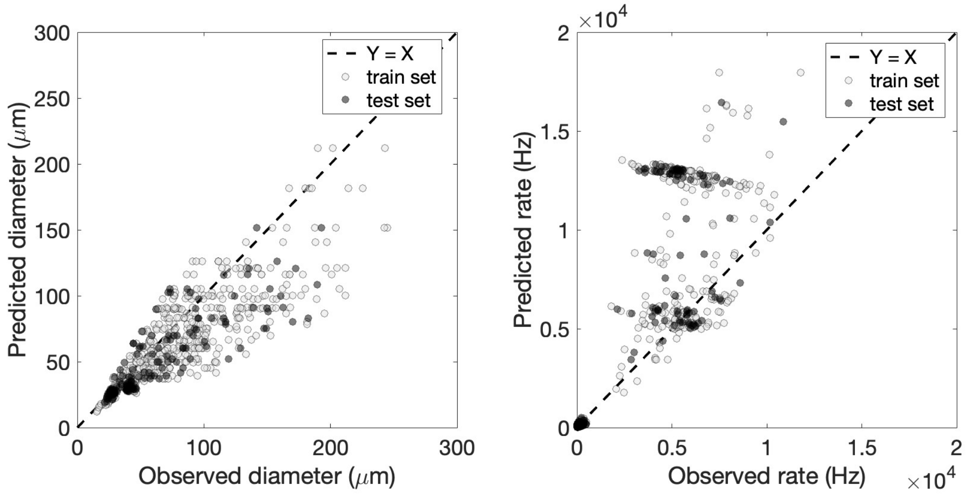


Supplementary Figure 9: The scaling law proposed by Ward et al.

### 2 Note 2: Effect of including viscosity ratio as an input of Neural Networks

Including viscosity ratio as the $8^{th}$ input of the neural networks minutely improved their accuracy for predicting droplet diameters and generation rates of the comprehensive dataset (MAPE of 7.15% vs 7.45%). However, including this parameter reduced its accuracy in predicting literature data (16.69% vs 12.4%). Additional statistical comparisons are provided in Supplementary Table 3.

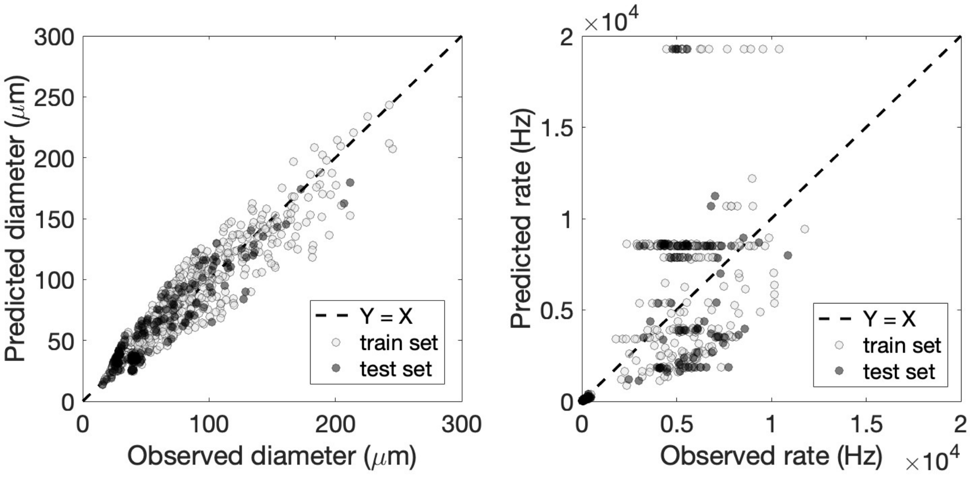


Supplementary Figure 10: The scaling law proposed by Zheya Liu et al.

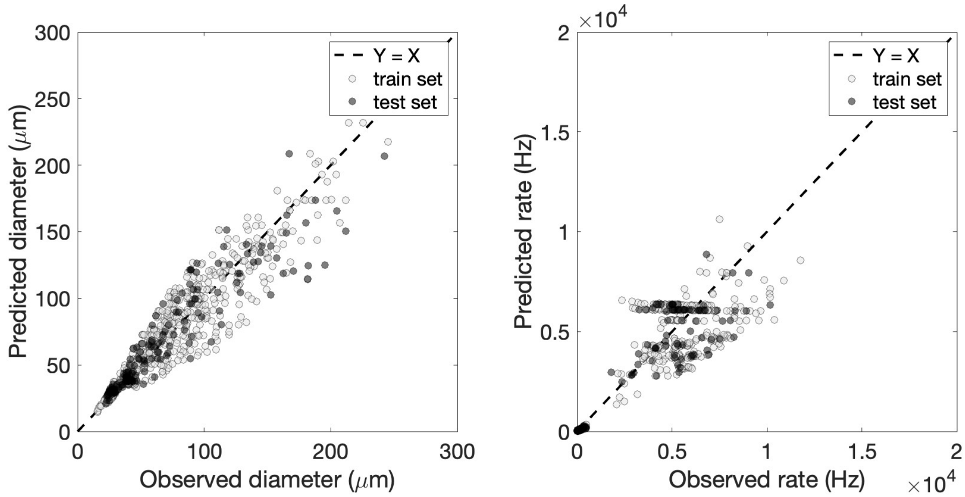


Supplementary Figure 11: The scaling law proposed by us.

### 3 Note 3: Percentage error relationship between droplet diameter and generation rate prediction

For both machine learning models, the MAPE for generation rate was approximately 3 times the MAPE for diameter. This is mathematically expected according to conservation of mass, as generation rate inversely scales with the $3^{rd}$ power of diameter.

$$F \sim \frac{1}{D^3} \tag{11}$$

For a small error in predicting diameter ($\delta_d$), using Taylor series expansion we have:

$$\frac{1}{(D+\delta_d)^3} \approx \frac{1}{D^3} - \frac{3\delta_d}{D^4} + \mathrm{O}\left(\delta_d^2\right). \tag{12}$$

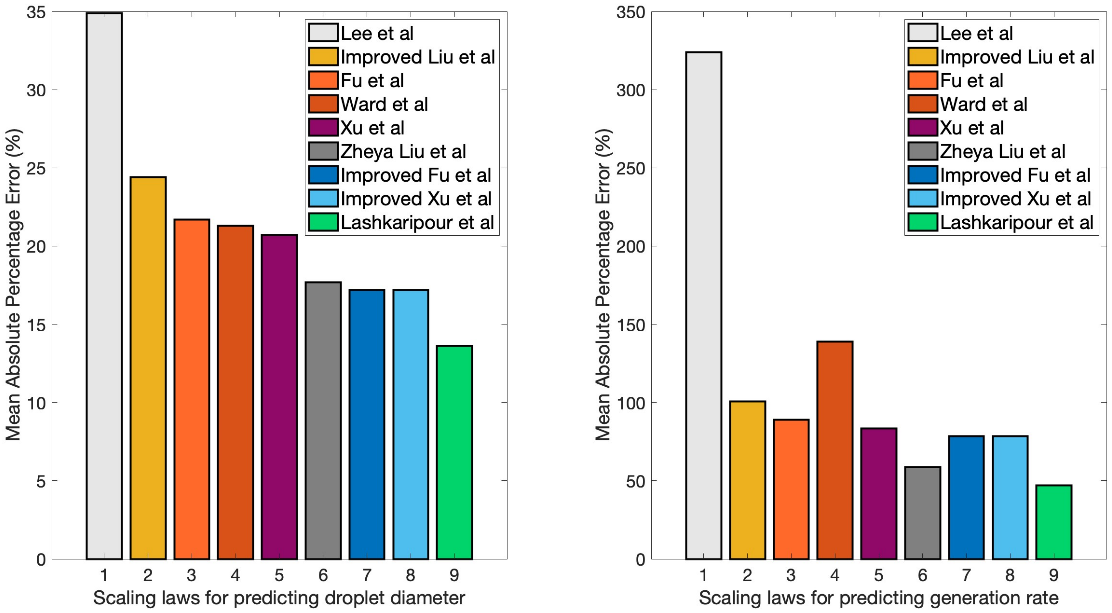


Supplementary Figure 12: Comparison of all the scaling laws in predicting droplet diameters and generation rates in our comprehensive dataset.

Therefore, prediction error for generation rate ($\delta_f$) is approximated by:

$$\delta_f \sim \frac{1}{D^3} - \frac{1}{(D+\delta_d)^3} \approx \frac{3\delta_d}{D^4} - \mathrm{O}\left(\delta_d^2\right). \tag{13}$$

For small errors of $\delta_d$, $\mathrm{O}\left(\delta_d^2\right)$ is negligible and $\delta_f$ is approximated by:

$$\delta_f \sim \frac{3\delta_d}{D^4}. \tag{14}$$

Percentage error in predicting generation rate can then be approximated by dividing Eq. 14 by Eq. 11. Therefore, for small values of $\delta_d$, percentage error for generation rate ($\varepsilon_f$) is approximated by 3 times the percentage error for diameter ($\varepsilon_d$):

$$\varepsilon_f = \frac{\delta_f}{F} \approx \frac{3\delta_d}{D} = 3\varepsilon_d \tag{15}$$

Therefore, we expect rate MAPE of accurate models to be 3 times the MAPE of the predicted diameter [3].

Supplementary Table 3: Effect of including viscosity ratio as the $8^{th}$ input parameter of the neural network on its accuracy for predicting droplet diameter and generation rate in the comprehensive dataset and their generalizability to predicting droplet diameters from previously published data.

| Parameter | MAPE | $R^2$ | MAE | RMSE |
|---|---|---|---|---|
| **Comprehensive dataset droplet diameter prediction** | | | | |
| Train set | 5.56 ±0.22% | 0.98 ±0 | 4.15 ±0.16$\mu$m | 6.70 ±0.21$\mu$m |
| Test set | 7.15 ±0.29% | 0.95 ±0 | 5.64 ±0.29$\mu$m | 9.36 ±0.43$\mu$m |
| **Comprehensive dataset generation rate prediction** | | | | |
| Train set | 16.73 ±0.34% | 0.97 ±0 | 248 ±12.07$Hz$ | 476.97 ±16.75$Hz$ |
| Test set | 21.91 ±0.93% | 0.97 ±0 | 262.84 ±12$Hz$ | 500.16 ±18.79$Hz$ |
| **Literature data diameter prediction** | | | | |
| Literature data | 16.69 ±1.07% | 0.82 ±0.03 | 12.19 ±1.02$\mu$m | 14.21 ±1.19$\mu$m |

Supplementary Table 4: DAFD 3.0 inputs and outputs for design automation of SEs of complete RPMI 1640 media with added 20% optiprep and 0.1% pluronic F-127 in dSurf HFE 7500, related to Fig. 6a of the main manuscript. The fluid properties of the dispersed and continuous fluids and the interfacial tension between them were inputted using the values provided in Table 1 of the main manuscript.

| **Desired performance** | | | **Proposed designs** | | | | | | |
|---|---|---|---|---|---|---|---|---|---|
| **Performance** | | **Constraints** | **Geometry ($\mu$m)** | | | | | **Flow rates ($\mu$l/hr)** | |
| Diameter | Rate | | Orifice | Depth | Outlet | Disp. inlet | Cont. inlet | Disp. fluid | Cont. fluid |
| 25 $\mu$m | 2000 Hz | geometry | 22.5 | 30 | 22.5 | 22.5 | 22.5 | 64.1 | 392.3 |
| 30 $\mu$m | 2000 Hz | geometry | 22.5 | 30 | 22.5 | 22.5 | 22.5 | 95.3 | 392.3 |
| 35 $\mu$m | 1000 Hz | geometry | 22.5 | 30 | 22.5 | 22.5 | 22.5 | 75.5 | 158.8 |

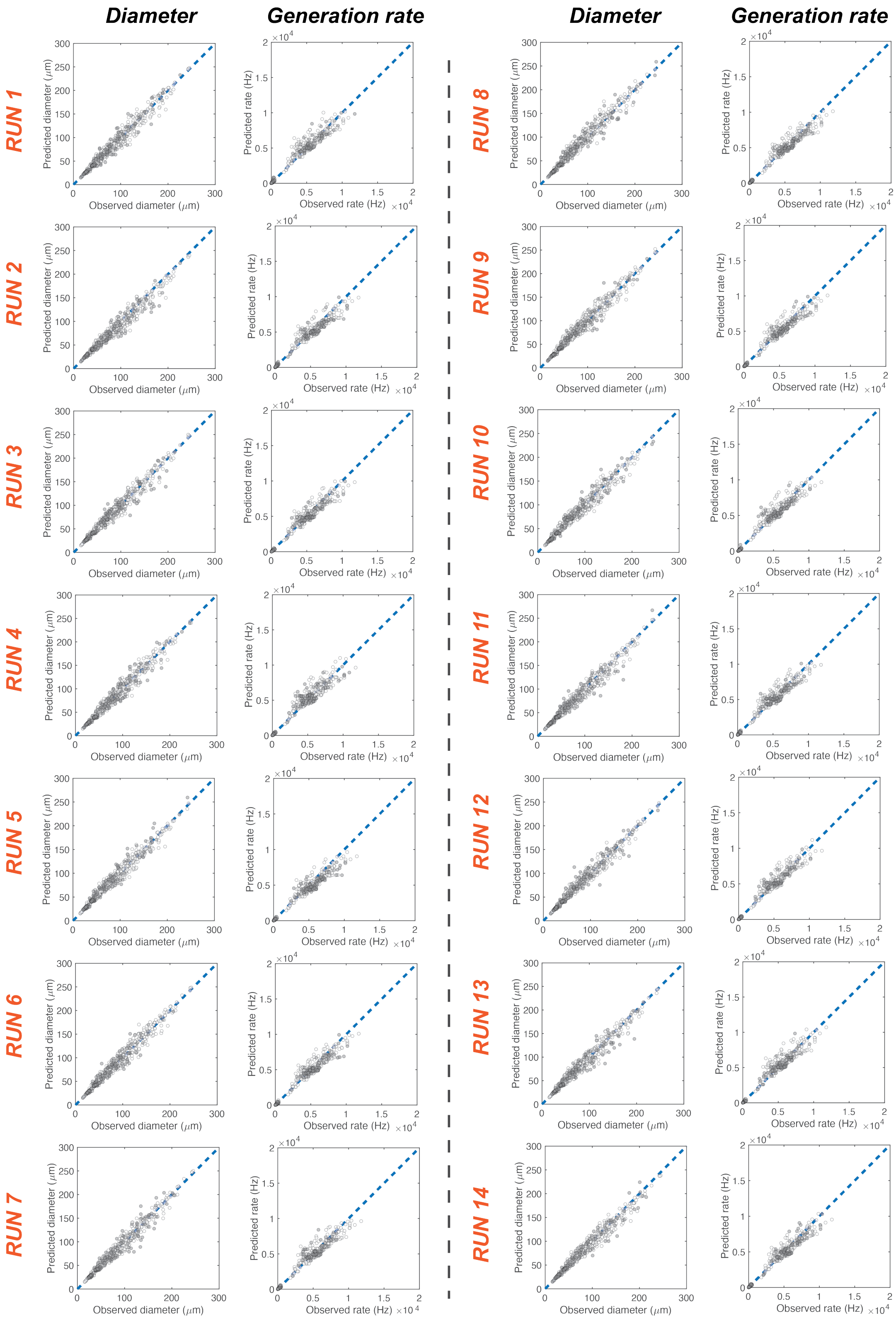


Supplementary Figure 13: Comparison of diameter and generation rate predictions of 14 random consecutive training runs of neural networks.

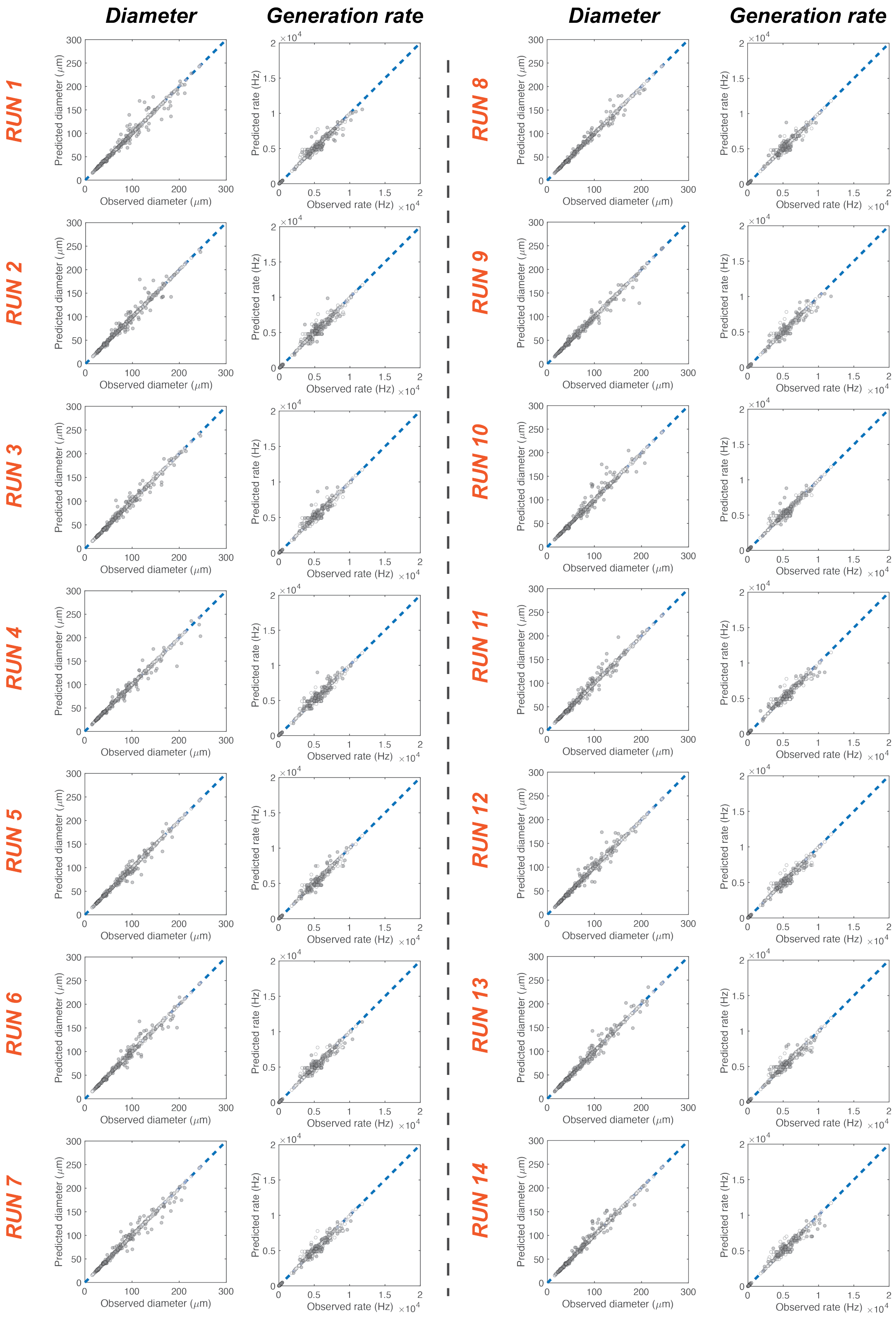


Supplementary Figure 14: Comparison of diameter and generation rate predictions of 14 random consecutive training runs of boosted decision trees.

Supplementary Table 5: DAFD 3.0 inputs and outputs for design automation of DEs using complete RPMI 1640 media with added 20% optiprep and 0.1% pluronic F-127, dSurf HFE 7500, and complete RPMI 1640 media with added 5% pluronic F-127 related to Fig. 6b of the main manuscript. The specified geometric constraints at FF2 were kept at 2x the same values at FF1 to facilitate stable DE generation. The fluid properties of the inner, middle, and outer fluids and the interfacial tensions between them were inputted using the values provided in Table 1 of the main manuscript.

| **Desired performance** | | | **Proposed designs** | | | | | | | |
|---|---|---|---|---|---|---|---|---|---|---|
| **Performance** | | **Constraints** | **Geometry at FF1 ($\mu$m)** | | | | | **Flow rates ($\mu$l/hr)** | | |
| Inner diam. | Outer diam. | | Orifice | Depth | Outlet | Disp. inlet | Cont. inlet | Inner | Middle | Outer |
| 25 $\mu$m | 50 $\mu$m | geometry | 22.5 | 30 | 22.5 | 22.5 | 22.5 | 100 | 650 | 4000 |
| 30 $\mu$m | 50 $\mu$m | geometry | 22.5 | 30 | 22.5 | 22.5 | 22.5 | 100 | 375 | 2500 |
| 35 $\mu$m | 50 $\mu$m | geometry | 22.5 | 30 | 22.5 | 22.5 | 22.5 | 125 | 200 | 2000 |
| 40 $\mu$m | 50 $\mu$m | geometry | 22.5 | 30 | 22.5 | 22.5 | 22.5 | 525 | 400 | 5500 |
| 25 $\mu$m | 55 $\mu$m | geometry | 22.5 | 30 | 22.5 | 22.5 | 22.5 | 50 | 500 | 2500 |
| 30 $\mu$m | 55 $\mu$m | geometry | 22.5 | 30 | 22.5 | 22.5 | 22.5 | 50 | 225 | 1500 |
| 35 $\mu$m | 55 $\mu$m | geometry | 22.5 | 30 | 22.5 | 22.5 | 22.5 | 125 | 200 | 2000 |
| 40 $\mu$m | 55 $\mu$m | geometry | 22.5 | 30 | 22.5 | 22.5 | 22.5 | 525 | 400 | 5500 |
| 30 $\mu$m | 45 $\mu$m | geometry | 22.5 | 30 | 22.5 | 22.5 | 22.5 | 400 | 1075 | 9500 |

Supplementary Table 6: Design parameter range and step size used for design automation of SE droplets.

| **Parameter** | **Parameter range & resolution** | | |
|---|---|---|---|
| | Minimum | Maximum | Step-size |
| Orifice width | 15 | 175 | 1.5 |
| Aspect ratio | 1 | 3 | 0.25 |
| Normalized oil inlet | 1 | 4 | 0.25 |
| Normalized water inlet | 1 | 4 | 0.25 |
| Expansion ratio | 1 | 6 | 0.25 |
| Flow rate ratio | 0.69 | 22 | 2 |
| Capillary number | 0.014 | 0.5 | 0.025 |

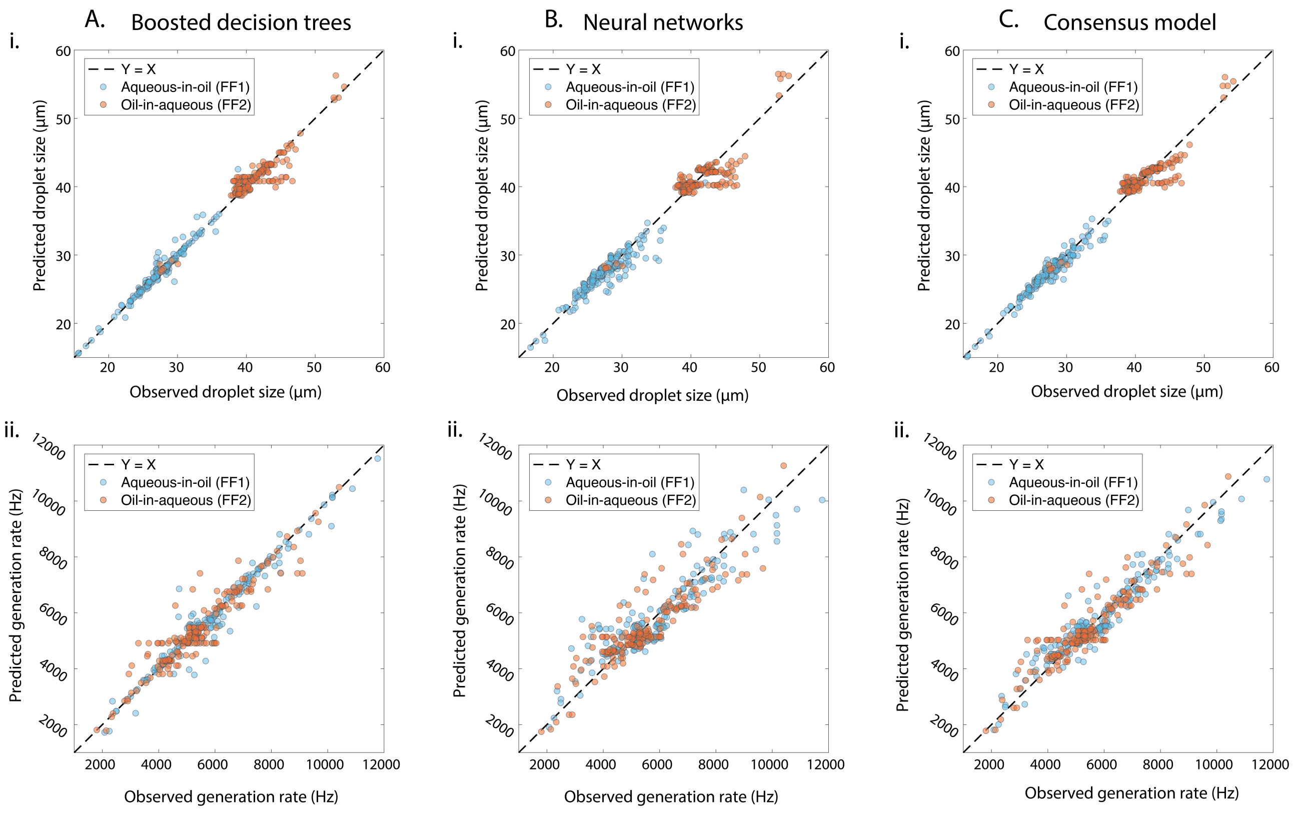


Supplementary Figure 15: Comparison of the trained machine learning models in predicting the inner and outer diameters of DEs in the comprehensive dataset.

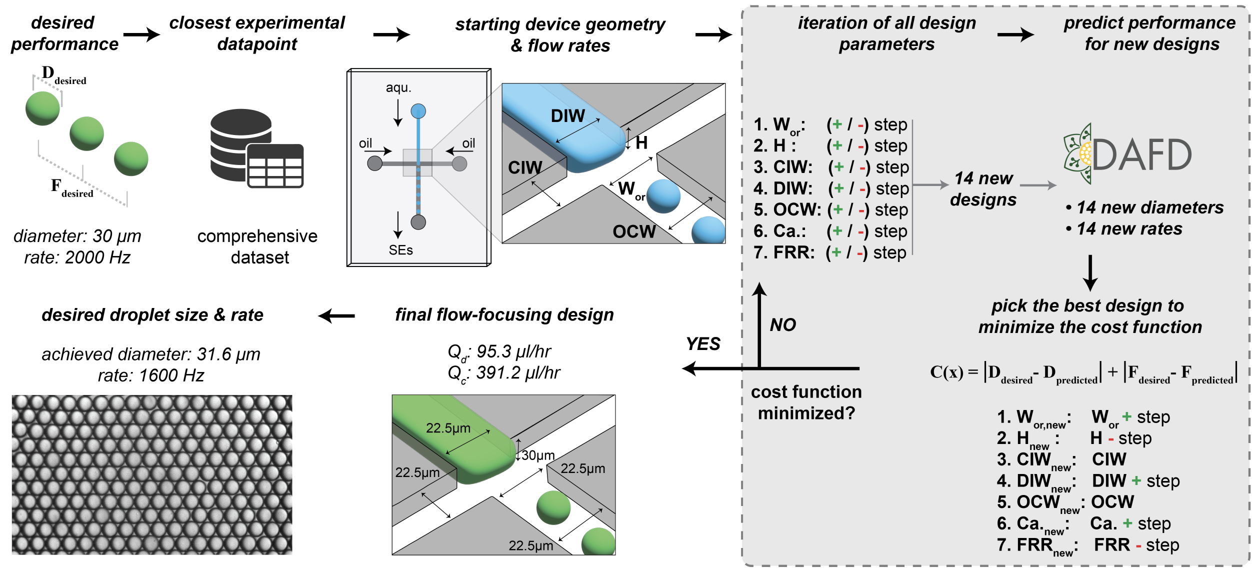


Supplementary Figure 16: The developed algorithm for design automation of single emulsions.

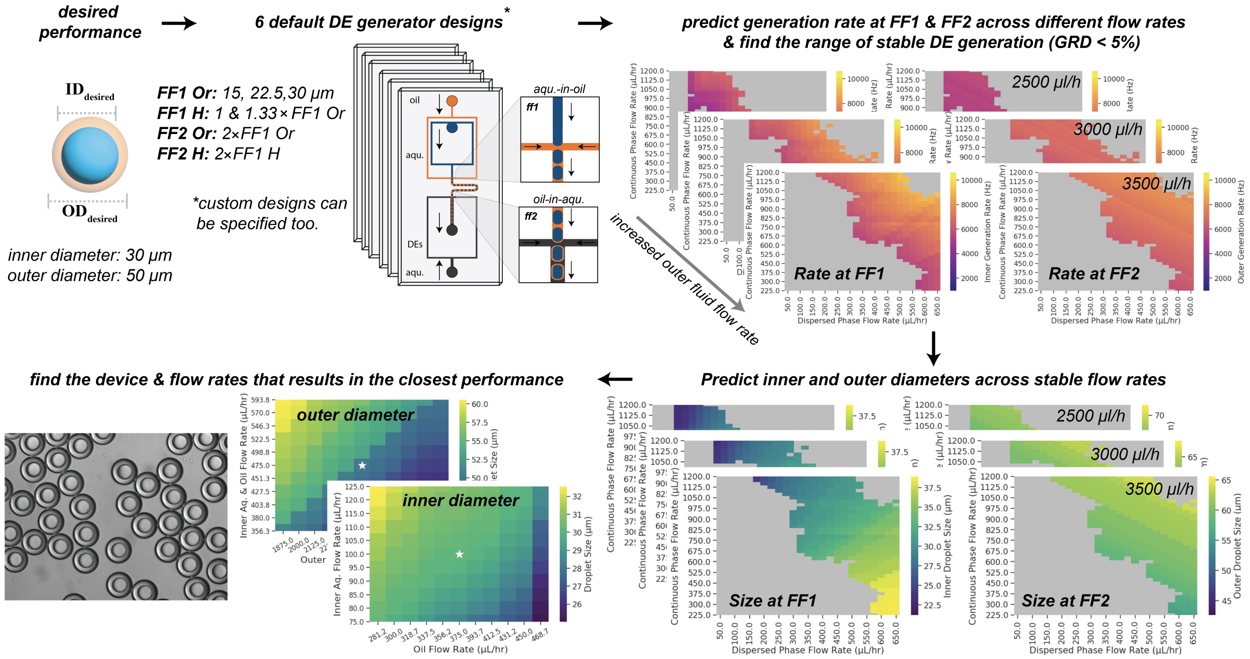


Supplementary Figure 17: The developed algorithm for design automation of double emulsions takes the desired inner and outer diameters and the fluid properties of the inner, middle, and outer fluids and find the device geometry and flow rates that results in the user-specified desired performance.

### Supplementary References

[1] Loïc Chagot, César Quilodrán-Casas, Maria Kalli, Nina M Kovalchuk, Mark JH Simmons, Omar K Matar, Rossella Arcucci, and Panagiota Angeli. Surfactant-laden droplet size prediction in a flow-focusing microchannel: a data-driven approach. *Lab on a Chip*, 22(20):3848–3859, 2022.

[2] Taotao Fu, Yining Wu, Youguang Ma, and Huai Z Li. Droplet formation and breakup dynamics in microfluidic flow-focusing devices: From dripping to jetting. *Chemical engineering science*, 84:207–217, 2012.

[3] Ali Lashkaripour, Christopher Rodriguez, Noushin Mehdipour, Rizki Mardian, David McIntyre, Luis Ortiz, Joshua Campbell, and Douglas Densmore. Machine learning enables design automation of microfluidic flow-focusing droplet generation. *Nature communications*, 12(1):25, 2021.

[4] Wingki Lee, Lynn M Walker, and Shelley L Anna. Role of geometry and fluid properties in droplet and thread formation processes in planar flow focusing. *Physics of Fluids*, 21(3):032103, 2009.

[5] Haihu Liu and Yonghao Zhang. Droplet formation in microfluidic cross-junctions. *Physics of Fluids*, 23(8):082101, 2011.

[6] Zheyu Liu, Maojie Chai, Xin Chen, Seyed Hossein Hejazi, and Yiqiang Li. Emulsification in a microfluidic flow-focusing device: Effect of the dispersed phase viscosity. *Fuel*, 283:119229, 2021.

[7] Thomas Ward, Magalie Faivre, Manouk Abkarian, and Howard A Stone. Microfluidic flow focusing: Drop size and scaling in pressure versus flow-rate-driven pumping. *Electrophoresis*, 26(19):3716–3724, 2005.

[8] Jian Hong Xu, SW Li, Jing Tan, and GS Luo. Correlations of droplet formation in t-junction microfluidic devices: from squeezing to dripping. *Microfluidics and Nanofluidics*, 5:711–717, 2008.
